# Molecular Characterization of Vaginal Microbiota Using a New 22-Species qRT-PCR Test to Achieve a Relative-abundance and Species-based Diagnosis of Bacterial Vaginosis

**DOI:** 10.1101/2024.03.18.585539

**Authors:** Ayodeji B. Oyenihi, Ronald Haines, Jason Trama, Sebastian Faro, Eli Mordechai, Martin E. Adelson, John Osei Sekyere

## Abstract

**Background:** Numerous bacteria are involved in the etiology of bacterial vaginosis (BV). Yet, current tests only focus on a select few. We therefore designed a new test targeting 22 BV-relevant species.

**Methods:** Using 946 stored vaginal samples, a new qPCR test that quantitatively identifies 22 bacterial species was designed. The distribution and relative abundance of each species, α- and β-diversities, correlation and species co-existence were determined per sample. A diagnostic index was modeled from the data, trained, and tested to classify samples into BV-positive, BV-negative, or transitional BV.

**Results:** The qPCR test identified all 22 targeted species with 95 – 100% sensitivity and specificity within 8 hours (from sample reception). Across most samples, *Lactobacillus iners, Lactobacillus crispatus, Lactobacillus jensenii, Gardnerella vaginalis, Fannyhessea (Atopobium) vaginae, Prevotella bivia,* and *Megasphaera sp. type 1* were relatively abundant. BVAB-1 was more abundant and distributed than BVAB-2 and BVAB-3. No *Mycoplasma genitalium* was found. Inter-sample similarity was very low, and correlations existed between key species, which were used to model, train, and test a diagnostic index: *MDL-BV index*. The *MDL-BV index*, using both species and relative abundance markers, classified samples into three vaginal microbiome states. Testing this index on our samples, 491 were BV-positive, 318 were BV-negative, and 137 were transitional BV. Although important differences in BV status were observed between different age groups, races, and pregnancy status, they were statistically insignificant.

**Conclusion:** Using a diverse and large number of vaginal samples from different races and age groups, including pregnant women, the new qRT-PCR test and *MDL-BV index* efficiently diagnosed BV within 8 hours (from sample reception), using 22 BV-associated species.

**Lay summary/Importance/Significance:** Bacterial vaginosis (BV) affects nearly 30% of women between 14 – 49 years old, increasing the risk and complications of endometriosis, pelvic inflammatory disease, pre-term births and low-birth weights, STIs, and cervicitis. Notwithstanding, BV’s diagnosis and etiology remain elusive. Not all BV-causing bacteria can be seen under a microscope or grown in a laboratory, making detection difficult. Moreover, current molecular BV diagnostic tests focus on a few signature species and thereby do not characterize the true state of the vaginal microbiome. Therefore, we designed a new Real-Time PCR test that identifies and quantifies 22 bacteria important in the prevention and development of BV. The data obtained from this test were further used to design and test a model that can easily use the relative abundance of the 22 species in any given vaginal sample to diagnose its BV status: all within 8 hours (from sample reception). The expansion of the bacterial spectrum in our new test enhances its resolution and broadens its diagnostic capacity to reduce false diagnoses and improve therapy.

## 1. Introduction

Bacterial Vaginosis (BV), a condition in which the normal lactobacillus-rich vaginal microbiome becomes dominated by polymicrobial anaerobic bacterial species under non-acidic pH, remains the most common cause of abnormal vaginal discharge in reproductive-age women, with an estimated prevalence rate of 29% in North America [1,2]. The normal vaginal microbiome is dominated by three major *Lactobacillus* species, *L. crispatus, L. jensenii,* and *L. gasseri,* which protect the vagina by producing lactic acid, hydrogen peroxide, and bacteriocins that suppress bacterial growth [3]. *Lactobacillus iners*, however, is quite enigmatic as it occurs in both healthy and unhealthy vaginal environments; *L. iners*, unlike *L. crispatus, L. jensenii,* and *L. gasseri,* produces the human non-functional L form of lactic acid [4].

Although BV is asymptomatic in many women, it is associated with the development of common obstetric and gynecologic complications [5–7]. It also increases the risk of contracting HIV and other sexually transmitted infections (STIs) and pelvic inflammatory disease (PID) [8]. Notably, BV recurrence after treatment is common (estimated to affect 50% of women annually [9–11]) and may be due to re-infections from sexual partners or failure of current treatment options [10,12]. Besides the health impacts of recurrence, BV treatment costs are increasing annually, specifically in the USA: this economic impact is particularly pronounced in BV-associated preterm births and other obstetric complications [13,14].

To diagnose BV, clinicians rely mostly on the classical clinical signs and symptoms outlined in Amsel’s criteria [15] or on the microscopically based Nugent score [16]. While these standard diagnostic methods have been effective over the years, they have been confronted with limitations such as subjectivity and the inability to accurately identify pathogens. The development of culture-independent molecular diagnostics has enabled the detection of non-cultivable bacterial species associated with BV and continues to revolutionize infectious disease diagnosis [11,17,18]. Molecular tests such as real-time polymerase chain reaction (RT-PCR), combined with the traditional clinical diagnostic criteria, have greatly improved the sensitivity and specificity of detecting BV pathogens [11,17,18]. They are also applicable in monitoring patient response to antibiotic therapy [1,19]. This approach has proven more useful for identifying patients at risk for recurrent BV [3,9,12,14,20,21].

The etiology and pathogenesis of BV are still not fully understood, making it important to study the involved microbes for timely diagnosis and treatment. Particularly, the interactions between species within the BV microbiome that cause pathologies, treatment failure and recurrence, or healing needs further interrogations to enable a species biomarker-based diagnosis of BV.

Besides next-generation sequencing (NGS)-based metagenomics that target all genomes, current PCR diagnostics for BV mainly focus on a smaller spectrum of bacterial species. Hence, to increase the resolution and diagnostic power of PCR-based BV diagnostics [22], we designed a new 22-species quantitative Real-Time PCR assay that quantitatively detects 22 vaginal bacterial species within 8 hours (from sample reception). This newly designed proprietary RT-PCR assay (Bacterial Vaginosis (with Lactobacillus Profiling) Panel^®^, Medical Diagnostic Laboratories, L.L.C. (MDL), New Jersey, USA) was used to screen 946 vaginal samples routinely obtained from different health centers across the United States. We further used the results, with assistance from machine-learning algorithms (Decision Trees and Random Forests) to design, train, and test an index (herein termed the *MDL-BV index*) that used a relative-abundance and species-based markers to classify the vaginal microbiome in three categories: BV-negative, transitional BV, and BV-positive.

## 2. Materials and Methods

### 2.1 Specimen collection and processing

Clinical vaginal samples are routinely obtained from different healthcare centers across the United States for diagnostic processing at MDL. Historical vaginal specimens (n = 946) marked for disposal were received in *One*Swab^®^ (Copan Diagnostics, CA, USA) or ThinPrep^®^ (Hologic, MA, USA) transport media in a Clinical Laboratory Improvement Amendments (CLIA)-certified infectious disease laboratory facility between January and June 2023 and stored at −80 °C were selected randomly for this study. This included specimens from symptomatic, asymptomatic, pregnant, or non-pregnant females and from whom vaginal profiling was requested. For each biological specimen that arrives at the laboratory facility, specimen accessioning occurs that assigns a random MDL number to ensure the de-identification of specimens. To further the de-identification of specimens during this study, samples that matched our collection criteria were randomized and the MDL numbers associated with each sample were not recorded.

### 2.2. Targeted bacterial species

The newly designed quantitative real-time PCR (qRT-PCR) assay (Bacterial Vaginosis (with Lactobacillus Profiling) Panel^®^ is a BV diagnostic assay designed by MDL to qualitatively and quantitatively detect 22 bacterial species that are found in the eubiotic and dysbiotic vaginal microbiome. These species are *L. crispatus, L. jensenii, L. gasseri, L. iners, L. acidophilia, Gardnerella vaginalis, Fannyhessea vaginae* (*Atopobium vaginae*)*, Megasphaera* sp. types 1 and 2, *Prevotella bivia*, Bacterial Vaginosis-Associated Bacterium (BVAB) 1-3, *Ureaplasma urealyticum, Mycoplasma hominis, Mycoplasma genitalium, Mobiluncus curtisii, Mobiluncus mulieris, Sneathia sanguinegens, Bifidobacterium breve, Bacteroides fragilis*, and *Streptococcus anginosus* [19,23–25].

### 2.3 DNA preparation

DNA from the vaginal specimens were extracted according to validated in-house laboratory protocols using QIAamp^®^ DNA Mini Kit (QIAGEN, Maryland, USA) and the X-tractor Gene^®^ DNA workstation (QIAGEN, Maryland, USA) with slight modifications. Canine Herpes Virus DNA was spiked into the samples and subsequently detected as an internal extraction control.

To serve as amplification standards, plasmids for each of the 22 bacterial species were generated by whole-gene synthesis (Genewiz^®^, Azenta Life Sciences, Waltham, USA). Briefly, a species-specific region of each species was amplified, synthesized, and cloned into the pUC-GW-AMP plasmid vector with an ampicillin-resistance marker. These plasmids were transformed into chemically competent *Escherichia coli* cells (One Shot™ TOP10, ThermoFisher Scientific, New Jersey, USA) and grown in liquid Lysogeny Broth (LB) containing 50 µg/mL Ampicillin. Plasmid DNA was isolated from overnight cultures using Wizard^®^ Plus SV Minipreps DNA Purification Systems (Promega, Wisconsin, USA) according to the manufacturer’s protocol. Extracted DNA was subsequently quantified spectrophotometrically using NanoDrop 1000 equipment (ThermoFisher Scientific, New Jersey, USA). Standard 10-fold serial dilutions of 10^8^ to 10^1^ DNA copies/µL were prepared for each bacterial species.

### 2.4 Identification of bacterial species by multiplex qRT-PCR

Each vaginal and plasmid control DNA sample were amplified in triplicates using species-specific primers and fluorescent probes on a CFX384 Touch Real-Time PCR Detection System (Bio-Rad, California, USA). The specific primer and probe sequences, as well as the multiplex RT-PCR conditions in the Bacterial Vaginosis (with Lactobacillus Profiling) Panel by Real-Time PCR assay are to MDL. An aliquot of 0.5μL of DNA was used as the template in a 4μL total PCR reaction mixture containing four primer sets, four probes, and the in-house—prepared mastermix of Taq DNA polymerase and dNTPs (MDL, New Jersey, USA) in multiplex PCR reactions. Human β-globulin DNA served as the internal control while no-template control was included to account for potential extraneous nucleic acid contamination. After amplification, a standard curve (fluorescence vs cycle number) was generated for each species and the target DNA copy for each sample was extrapolated. A 10^2^ copies/μL concentration in each sample was established as the positive cutoff for a given bacterium. The relative percentage concentration of all the bacterial species identified in each vaginal specimen was then computed and tabulated (Table S1).

### 2.5 Relative abundance- and species-based classification of BV

The mean, median, standard deviation, and concentration distribution of each species across samples as well as alpha-diversity, Shannon index, beta-diversity (Bray-Curtis Dissimilarity matrix), and relative abundance were calculated from the gDNA concentration data (obtained from the qPCR amplification results of each species from the samples) and represented as box and Whisker plots, histograms, heatmaps, and charts. Using the distribution patterns in the data, a relative abundance or concentration-based cut-off criteria was established and used to define BV-positive, BV-negative, and transitional BV status. Using the species-species correlation and co-existence data as well as previous research from literature, a species-based criteria was also established for determining BV-positive, BV-negative, and transitional BV status. A two-tier system was then created by synthesizing the species-based and concentration-based BV diagnosis criteria, which was used to interpret the PCR results. This criterion was called the *MDL-BV index*, which was further trained and tested on the qPCR and demographics data using machine-learning algorithms (Decision Trees and Random Forests).

### 2.6 Statistical analysis

Statistical analyses were performed with Prism version 10.1.2 (Graph-Pad, California USA) or the R statistical package. Quantitative data obtained in this study were expressed as the meanLJ±LJstandard error of the mean (SEM). To detect bacterial associations and correlations, Spearman’s rank correlation coefficient was chosen. The data were parsed through Python (and Biopython) to calculate the significance of the coexistence between any two species using Chi-square and T-test.

Three types of groupings were used in this (Chi-square and T-test) analysis: (1) all *Lactobacillus* species were grouped into one and their absence/presence were compared with the absence/presence of each non-*Lactobacillus* species; (2) all *Lactobacillus* species except *L. iners* were grouped into one and their absence/presence were compared with the absence/presence of each non-*Lactobacillus* species; (3) the presence/absence of each *Lactobacillus* species was compared with that of each non-*Lactobacillus* species. The resulting data were tabulated in Table S2, and the significant results were filtered out and used to generate heat maps. The results were only considered significant if *P* <LJ0.05.

## 3. Results

### 3.1 Demographics

The vaginal samples were obtained from 946 women whose average age was 34.8 years, with a standard deviation of 13.3 years. The ages ranged from 18 to 84 years. The median age was 32 years; hence, the age distribution is somewhat skewed towards younger women (Figure 1). Fifty-three women reported as pregnant while the rest were either not pregnant or unknown. Race data was obtained for 245 samples: White (n = 145 samples), Black (n = 79), “Other race” (n = 15), Asian (n = 4), and Native American (n = 2) (Table S1). The “Other race” category refers to those not falling within any of the four races above.

**Figure 1.**
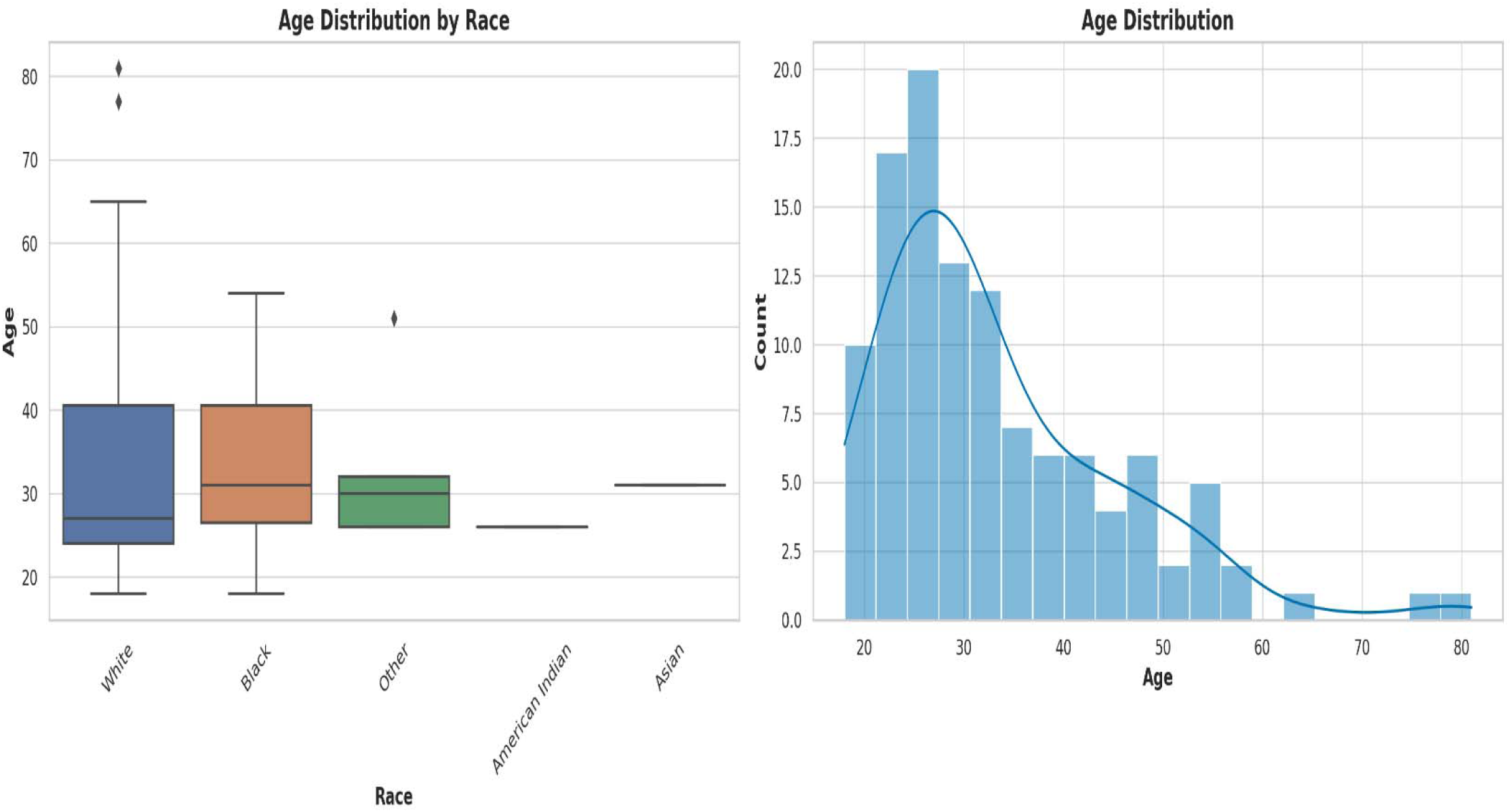
Box plot and histogram showing the age distribution among the races from which the vaginal samples were obtained. The box plot on the left, showing the age distribution among Whites, Blacks, Other (races), Native American or Alaska Natives, and Asians. Whites were the largest population, followed by Blacks. Other races are those who belong to none of the four races. The median ages among the races were very close within 28 and 35. An age distribution histogram on the right, shows the overall age distribution of participants in the study. The histogram includes a kernel density estimate (KDE) to show the smooth distribution of ages. The data is somewhat right-skewed, indicating a younger population with fewer older women. Most women fell into the 20-40 age range, with a peak around the late 20s to early 30s.

The age distribution of the races (Fig. 1) shows a broader distribution of ages within the White population (with outliers) than that of the other races, albeit the median ages across all races fell within a narrow range of 28 – 35 years. The age distribution of Whites ranged from 18 – 83 years, with 50% falling within 28 – 40 years. Whereas the ages of Blacks ranged from 18 to 55 years, 50% were within the same 28 – 40 years range. The “Other races”, which includes Hispanics/Latinos, had ages between 28 and 32 years and the median age was different among all the races (Fig. 1).

### 3.2 Identification and distribution of species across samples

Compared with the plasmid controls, the test was able to efficiently identify all of the targeted 22 species with 95 – 100% sensitivity and specificity. Except for M. *genitalium* that was not detected in any sample, all the other species were present in at least one of the 946 samples. The percentage of the count of the identified species across all the samples are as follows: *L. iners* (75%), *G. vaginalis* (65%), *F. vaginae* (53%), *Megasphaera* sp. type 1 (44%), and *P. bivia* (41%) (Figure 1). These are followed by *L. crispatus* (29%), *L. jensenii* (23%), BVAB1 (17%), *U. urealyticum* (13%), *L. gasseri* (12%), *M. hominis* (12%), *M. curtisii* (11%), and BVAB3 (10%). The remaining species had less than 10% relative abundance across all samples combined (Fig. 2).

**Figure 2.**
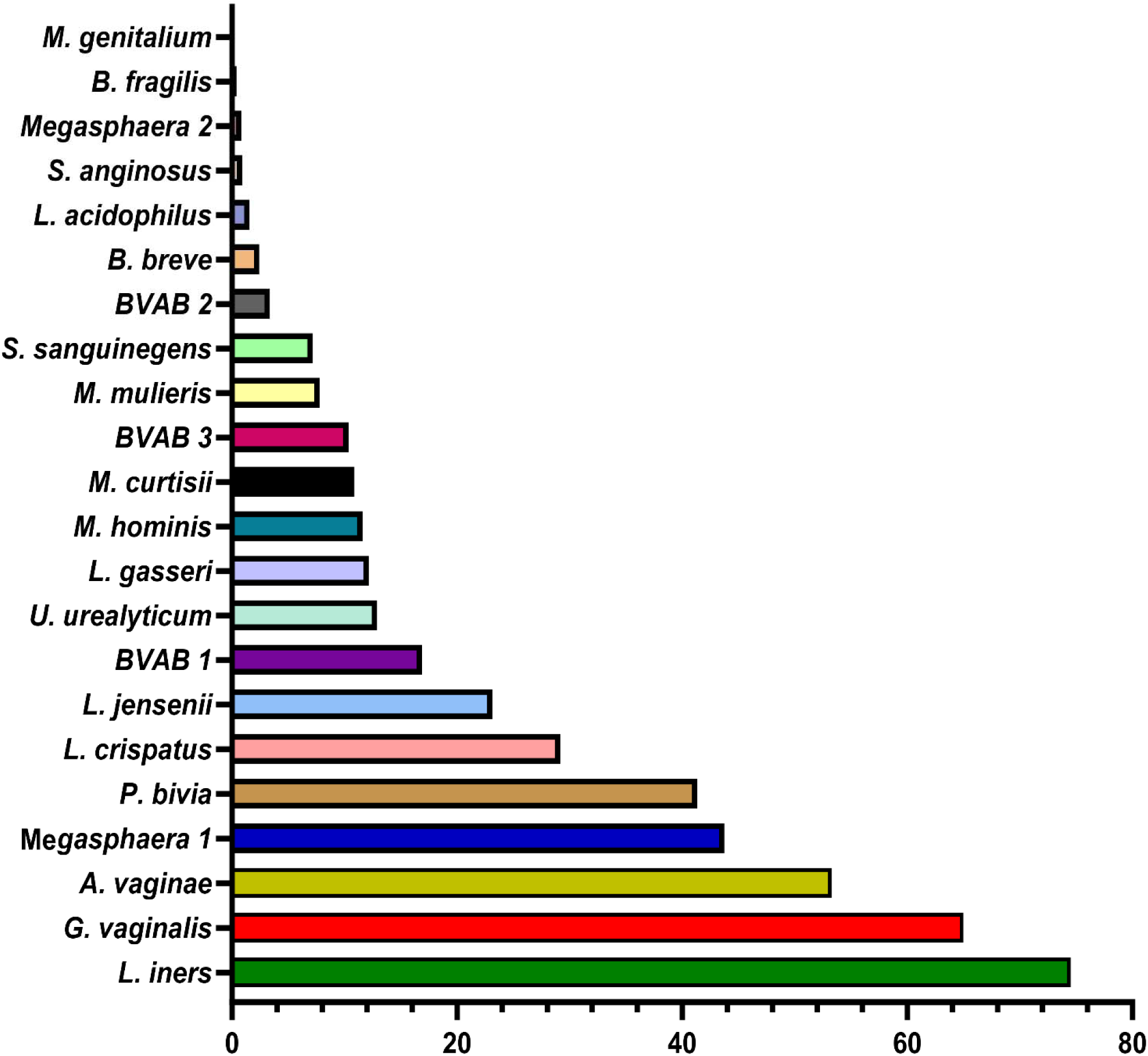
Percentage of the count of each species across all vaginal specimens (n = 946) as detected by qPCR. Among the 946 samples, it is observed that the most dominant species are *Lactobacillus iners, Gardnerella vaginalis, Atopobium (Fannyhessea) vaginae, Megasphaera sp. type 1, Prevotella bivia, Lactobacillus crispatus, and Lactobacillus jensenii.* The horizontal axis is the percentage of each species across all samples. No *Mycoplasma genitalium* was detected in any of all the samples.

To understand the distribution (spread) of the species and their concentrations across the samples, we used bar charts, histograms, and box and Whisker plots. *L. iners,* followed by *G. vaginalis, A. vaginae, Megasphaera sp.* type 1*, P. bivia, and L. crispatus* had better concentration distributions across the samples than the other species, with their median concentrations (10^2^ and 10^6^ gDNA copies/µL) being found in more than 100 samples. Indeed, most species had very low concentrations across all the samples (Fig. S1-S22). The relative concentration for each identified species ranged between 10^2^ and 10^8^ genomic DNA copies/µL and the median range was between 10^2^ and 10^6^ copies/µL (Fig. 3).

**Figure 3.**
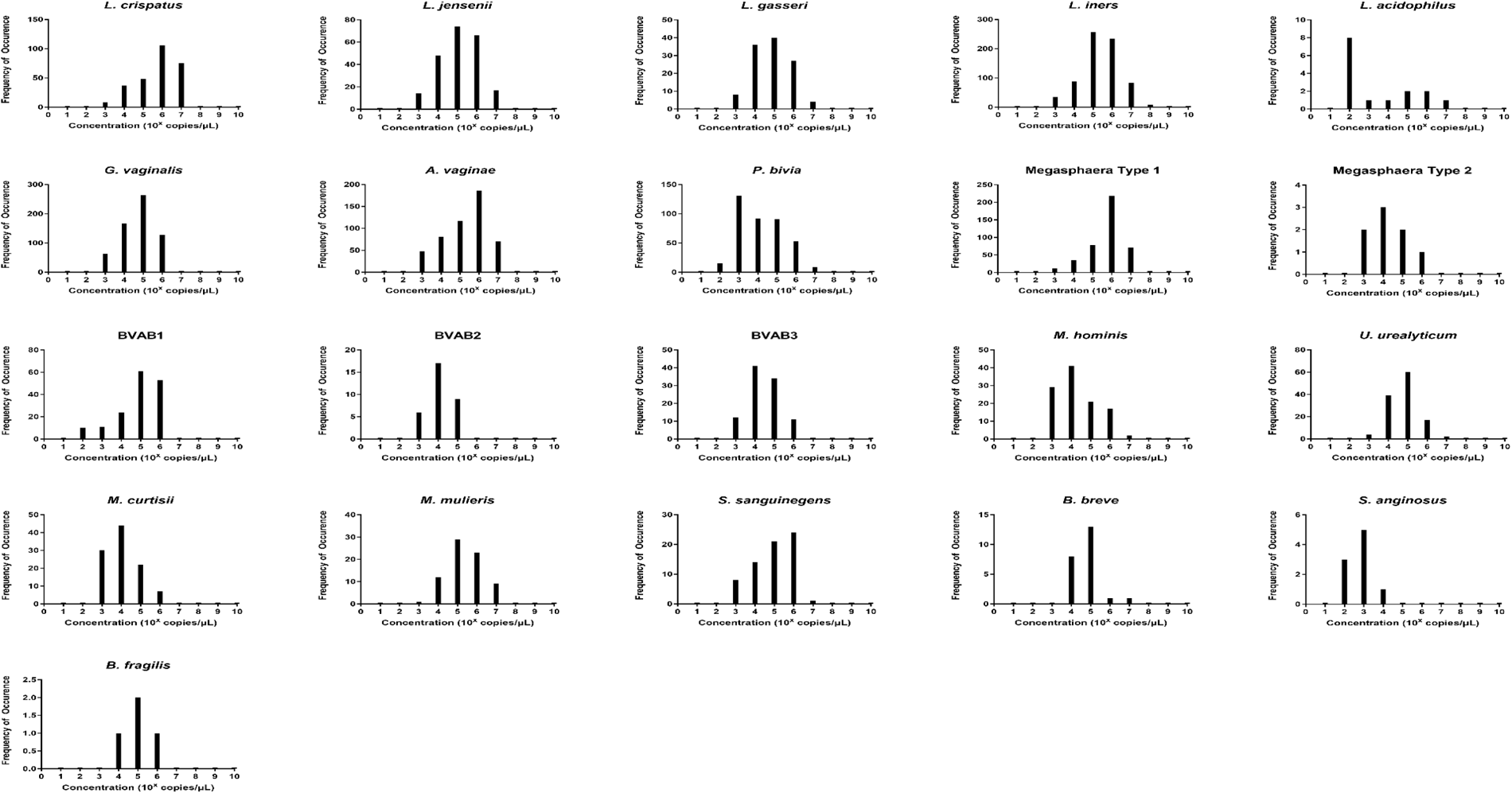
The relative concentration distribution of the identified bacterial species in vaginal swabs (n = 946). Each chart represents the concentration distribution per species; *M. genitalium* is not shown as was not detected in any of the samples. The vertical axis shows the count (frequency) of samples while the concentrations are shown on the horizontal axis. The median concentrations (10^2^ and 10^6^ gDNA copies/µL) of *Lactobacillus iners* and *L. crispatus, Gardnerella vaginalis, Atopobium (Fannyhessea) vaginae, Prevotella bivia,* and *Megasphaera sp. type 1* were found in more than 100 samples. The charts also do not show the count of samples with zero concentrations but only counts of samples with higher concentrations.

The concentrations of the *Lactobacillus* species detected in this study were in the order, *L. iners > L. crispatus > L. jensenii > L. gasseri > L. acidophilus* (Fig. 2). Additionally, *F. vaginae* and *G. vaginalis* occurred in similar amounts (∼10^5^ copies/µL) and were slightly more than the values obtained for *P. bivia* (10^4^ copies/µL). BVAB 1 was the most predominant of the BVABs, and its relative concentration was 44% and 31% more than BVAB3 and BVAB2, respectively. Notably, over 98% of the *Megasphaera* species identified in this study belonged to type 1 with a mean concentration of ∼10^6^ copies/µL. Of the Mycoplasmas, *U. urealyticum* was found in higher concentrations than *M. hominis*. Although *M. curtisii* was 39% more abundant, it occurred in lower concentrations in vaginal swabs than *M. mulieris*.

### 3.3 Relative abundance, *α*- and *β*-diversities

We further investigated the per-sample species relative abundance and richness as well as inter-sample species diversity using alpha and beta diversities. The relative abundance, shown as a heatmap and a box plot (Fig. 4), represents the abundance of each species across samples, normalized relative to the total concentration of microbial species within each sample. The relative abundance of the species perfectly mirrored their concentration distribution across the samples (Fig. 3; Fig. S1-S22), with *L. iners, L. jensenii, P. bivia, G. vaginalis, Megasphaera sp.* type 1, and *A. vaginae* being visibly abundant in many samples than the other species. However, the other less abundant species also presented greater variability in relative abundance across the samples, as shown in the outliers (black stars) and absence of boxes (Fig. 4B).

**Figure 4.**
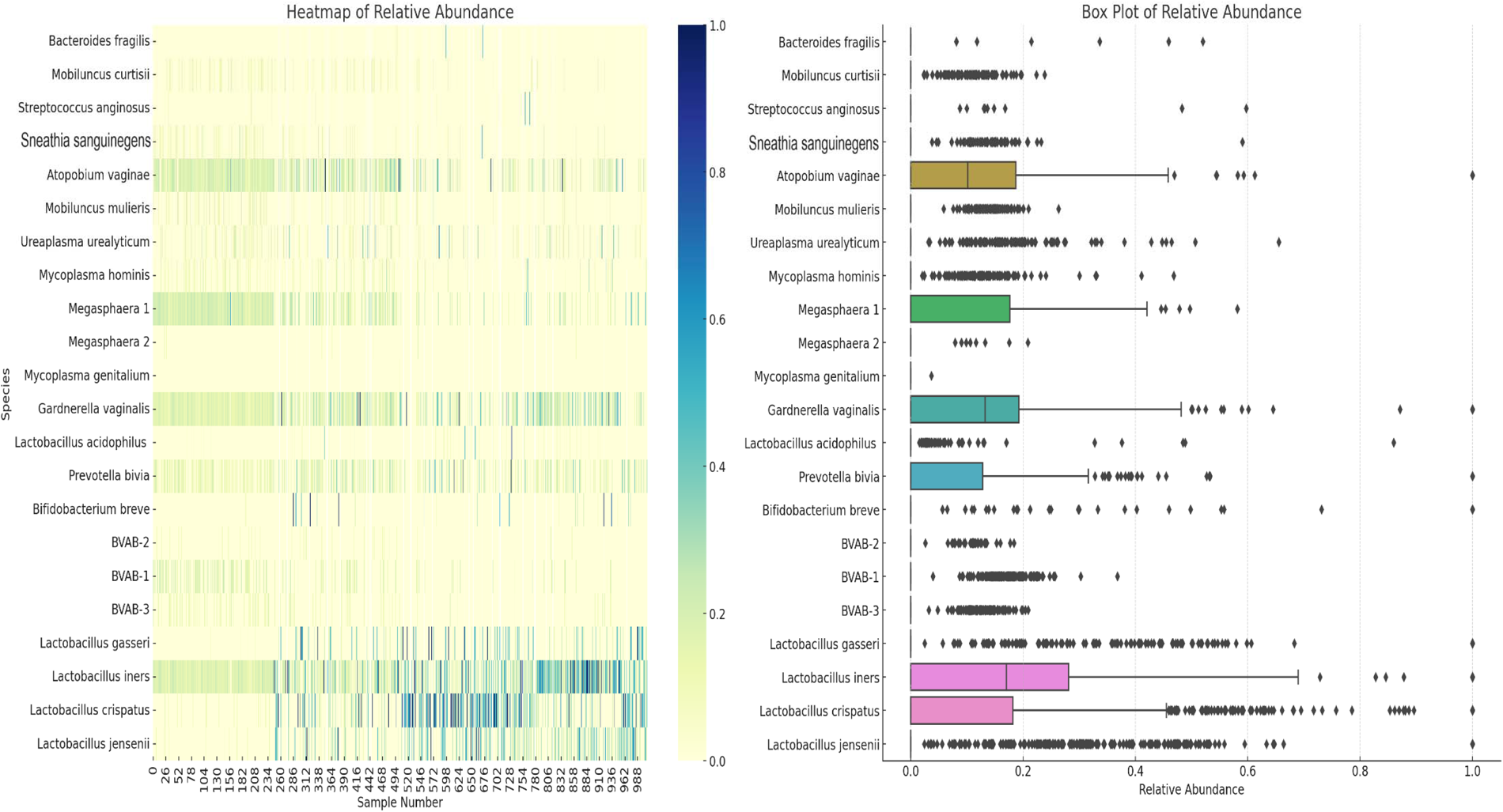
Relative abundance of the 22 species across all the 946 vaginal samples. The **heatmap** (**A**) highlights the distribution and intensity of species’ relative abundance across all samples, offering a color-coded representation that easily identifies the most prevalent species. The **box plot** (**B**) provides a statistical summary of each species’ relative abundance, including the median, interquartile range, and any outliers, offering insight into the variability and distribution of abundance for each species. This plot provides insights into the central tendency, spread, and outliers for the relative abundance of each microbial species within the dataset. Median values are represented by the line within each box. Interquartile range (IQR), indicating the middle 50% of the data, is shown by the box itself. Whisker extend to show the range of the data, i.e., 1.5 * IQR from the quartiles. Outliers are points outside the Whisker and are indicated as individual points.

The alpha diversity of the species within each sample was determined using species richness and Shannon diversity index (Fig. 5). The species richness shows that 50% of the samples had 2 – 6 species, with 8 – 12 species occurring in 100 – 300 samples (Fig. 5A and 5D). The Shannon index, which ranges from 0 (no diversity) to 5 (practically 3.5, i.e., most diverse), showed that 50% of the samples’ diversity index was between 0.7 – 1.8, with 200 – 280 samples having a diversity of > 2.0 (Fig. 4B and 4D).

**Figure 5.**
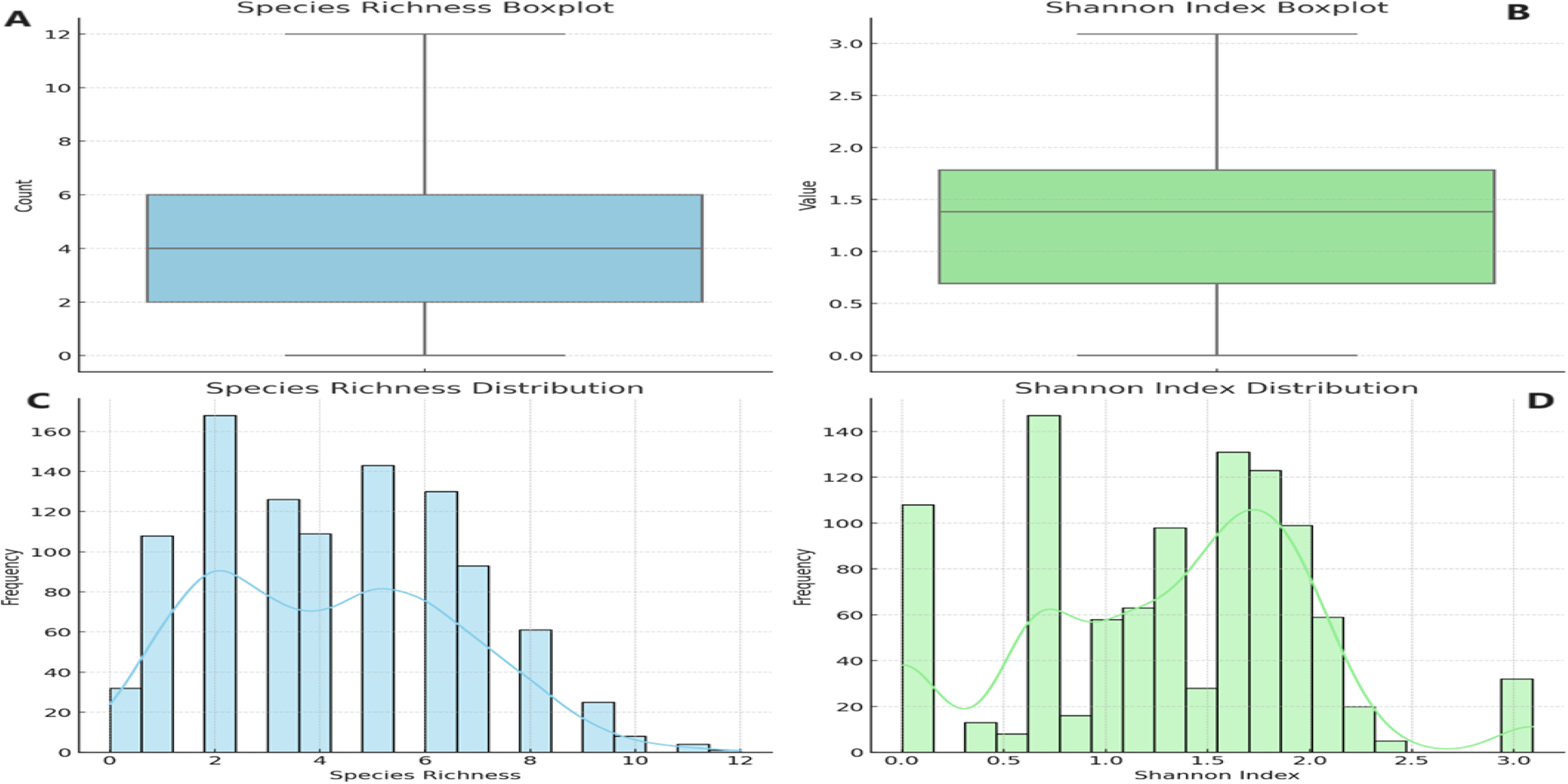
Alpha diversity of the 946 vaginal samples: species richness and Shannon index. Boxplot of Species Richness (**A**), showing the count of species present across samples. This boxplot provides a clear visualization of the central tendency and variability in species richness. Boxplot of the Shannon Index (**B**), indicating the diversity value that considers both abundance and evenness of species. Histogram of Species Richness Distribution (**C**), providing a view of the frequency distribution of species counts across samples. Histogram of Shannon Index (**D**) visualizes the distribution of the Shannon index across samples, reflecting both the abundance and evenness of species. The Shannon index is a more comprehensive measure of diversity, considering not just the presence of species but also their relative abundances.

Inter-sample (β-) diversity, using the Bray-Curtis dissimilarity index and associated principal component analysis (PCoA), showed that a few of the samples shared strong similarities to each other (Fig. 6). Samples with close similarity in species diversity are shown as blue (purple) while those with little similarity are shown as yellow. The abundance of yellow in the matrix shows how different most of the samples are from each other (Fig. 6A). The PCoA chart also reflected this, with a few samples clustering together while most of the samples were spatially separated from each other (Fig. 6B).

**Figure 6.**
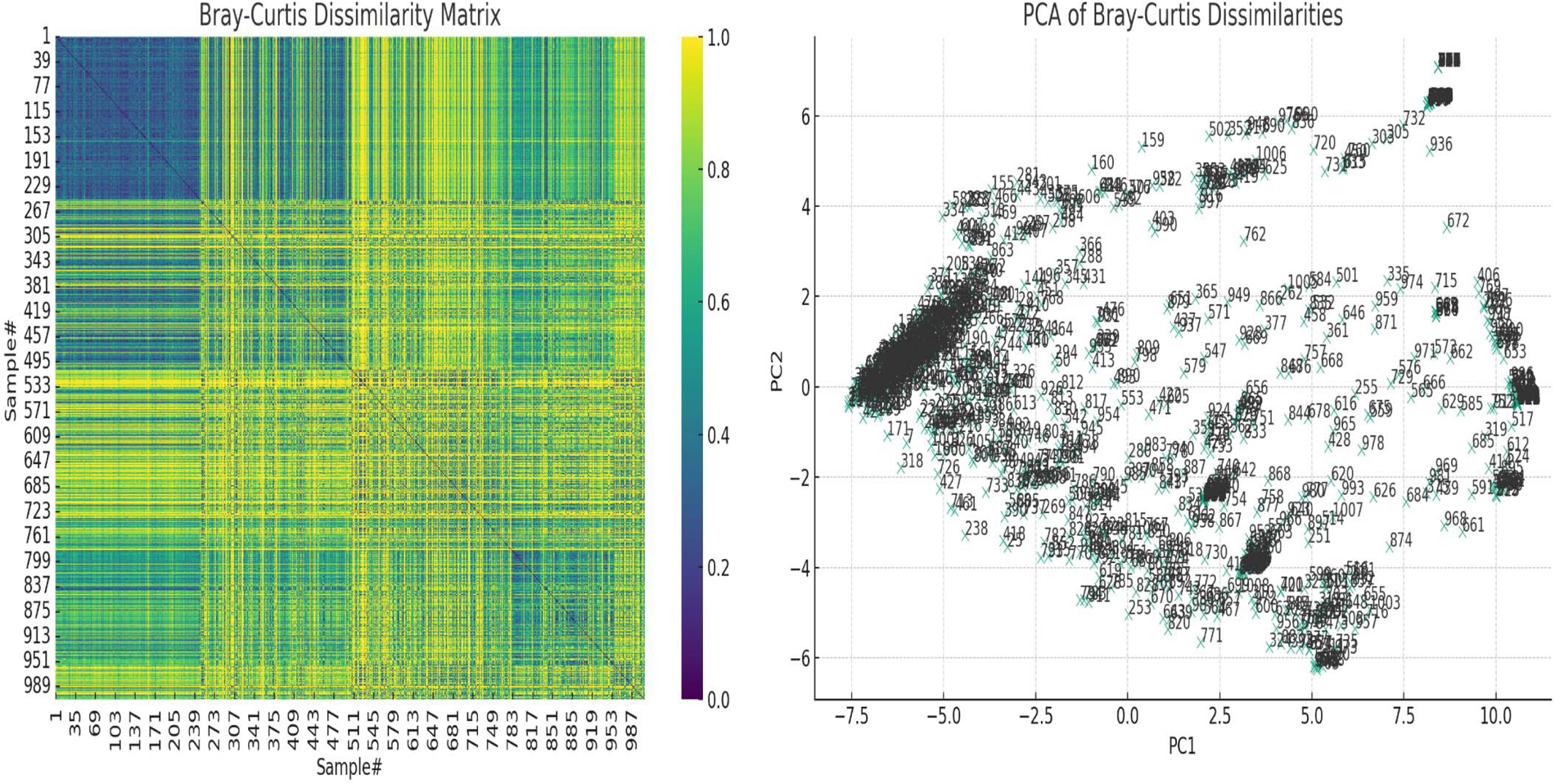
The Bray-Curtis dissimilarity matrix heatmap and associated principal component analysis (PCoA) for beta diversity. On the left, the Bray-Curtis Dissimilarity Matrix heatmap shows the pairwise dissimilarities between the samples. Higher values (closer to yellow, 1.0) indicate greater dissimilarity between samples, while lower values (closer to purple, 0.0) indicate greater similarity. On the right, the PCoA of Bray-Curtis Dissimilarities scatter plot visualizes the samples in a reduced two-dimensional space based on their dissimilarities. Each point represents a sample, and their positions reflect patterns of variation across the samples. Labels on the plot correspond to the sample numbers, helping to identify specific samples within the context of the PCoA. These show how dissimilar the samples were from each other.

### 3.4 Correlation and co-existence of species

We undertook a correlation, Chi-square and T-test analyses of the data to determine significance of the species-species co-existence in the samples (Table S2). We filtered out non-significant results and used the significant results to generate a heat map for easy visualization of the data (Fig. 7). Whereas the T-test provided a significant association between the presence/absence of the *Lactobacillus* species (as a group or without *L. iners*) and all the non-*Lactobacillus* species (BVAB-1 was significantly affected by the presence/absence of *L. iners* only but not by the other *Lactobacillus* species), the Chi-square test did not. Hence, the Chi-square test provided a more stringent cut-off than the T-test: individually, the *Lactobacillus* species were significantly associated with 14 species. In the absence of *L. iners,* the *Lactobacillus* species were not significantly associated with *B. fragilis* (which was only significantly associated with *L. gasseri*) but rather, with *Megasphaera* sp. type 2 (Fig.7).

**Figure 7.**
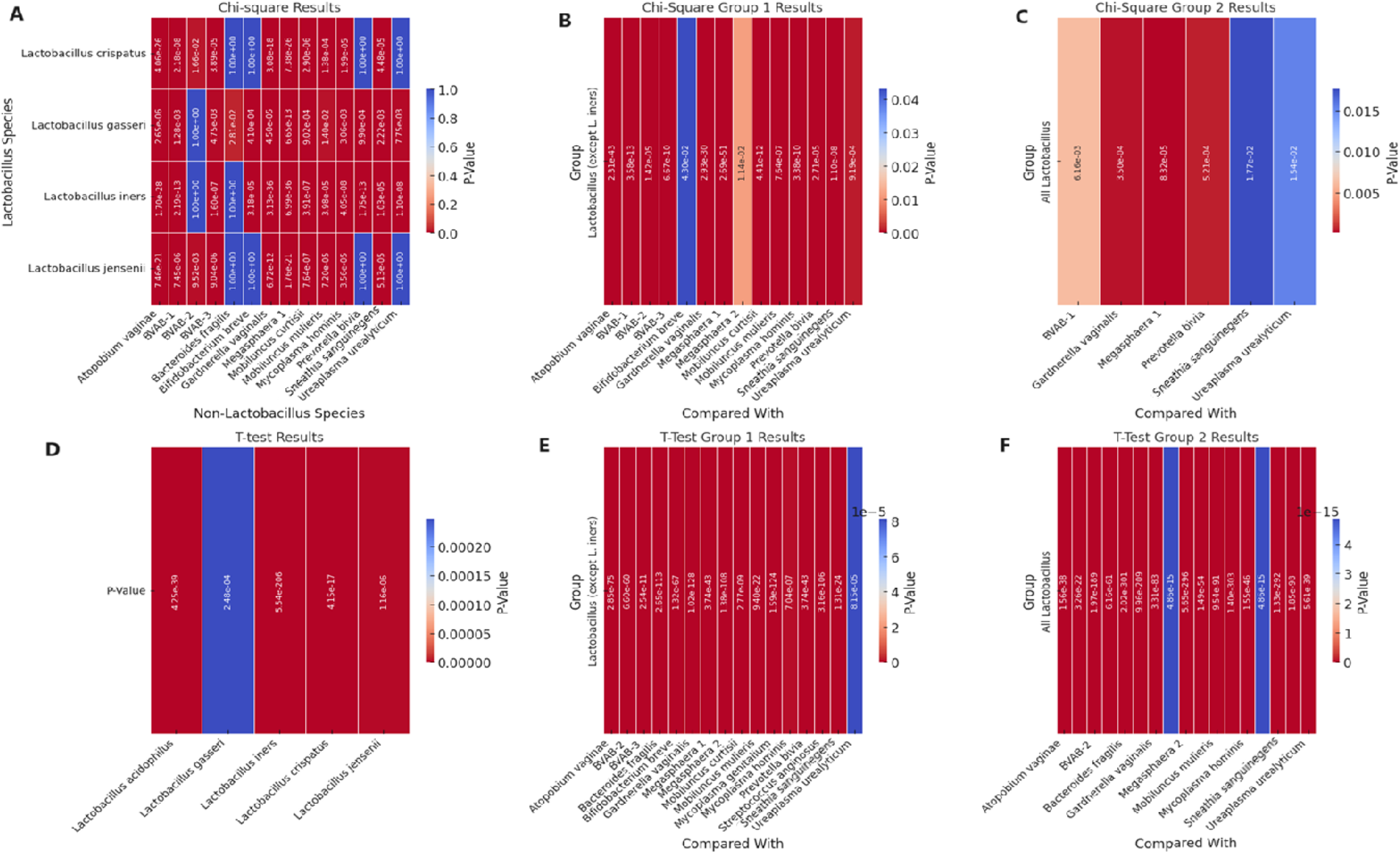
Heatmap of significant p-square values obtained from Chi-square and T-test analysis of pairwise comparisons between Lactobacillus and other non-lactobacillus species within the samples. **A** visualizes the significant Chi-square p-values between Lactobacillus species and non-Lactobacillus species. **B** (Group 1) visualizes the significant Chi-square p-values of all Lactobacillus species (except *L. iners*) against each non-Lactobacillus species. **C** visualizes the significant Chi-square p-values from grouped comparisons of all Lactobacillus species against each non-Lactobacillus species. **D** shows the significant T-test p-values of all Lactobacillus species against all the other non-Lactobacillus species grouped together (T-test could not get individual species comparisons because of the structure of the data). **E** visualizes the significant T-test p-values from T-Test Group 1, which involves grouped comparisons of all Lactobacillus species (except *L. iners*) against each non-Lactobacillus species. **F** visualizes the significant T-test p-values from the "T-Test Group 2" sheet, which involves grouped comparisons of all Lactobacillus species against each non-Lactobacillus species. Rows represent different Lactobacillus species. Columns represent different non-Lactobacillus species. The color intensity indicates the level of significance, with cooler colors (towards blue) indicating lower p-values (higher significance) and warmer colors (towards red) indicating higher p-values (lower significance). Cells filled with a p-value of 1.0 represent non-significant results that were not included in the original significant results table and are shown for completeness.

Therefore, the presence of BVAB-1 and *B. fragilis* were independent of *L. iners.* Furthermore, *L. crispatus* and *L. jensenii* were significantly associated with the presence/absence of the same non-*Lactobacillus* species except *B. fragilis, B. breve, P. bivia,* and *U. urealyticum* (Fig. 7). Although *L. acidophilus* was included in the Chi-square pairwise association test, it did not yield any significant results with any of the species (Table S2).

Using the Pearson’s correlation co-efficient, a clear pattern was observed regarding the co-existence of the 21 species with each other within the vaginal microbiota: all the *Lactobacillus species,* except *L. iners,* inversely correlated with most of the other non-*Lactobacillus* species while species such as BVAB (1-3), *Megasphaera sp.* type 1*, P. bivia, G. vaginalis, A. vaginae, M. hominis, M. mulieris, M. curtisii*, and *S. sanguinegens* were positively correlated with each other. Notably, there were species that had very little or no inverse correlation with the *Lactobacillus* sp. (except *L. iners*): *S. anginosus, B. breve, U. urealyticum, M. hominis, Megasphaera sp.* type 2, and *B. fragilis. S. anginosus, B. breve,* and *B. fragilis* were the only non-*Lactobacillus* species with an inverse correlation with *L. iners;* the other species had a positive correlation with *L. iners.* The BVAB species did not correlate with each other as BVAB -2 was less correlated with both BVAB-1 and BVAB-3; these latter two species, however, had a strong positive correlation (Fig. S23).

### 3.5 Relative abundance- and species-based diagnostic criteria

All the *Lactobacillus* sp. were bundled together as a marker of a normal vaginal microbiome (Group 1). Owing to the absence of a Nugent score or Amsel data for the samples, we used a species-based criteria to select species that are not found in normal vaginal flora: BVAB-1, -2, - 3, *B. fragilis,* and *S. anginosus.* These five species were bundled into a species marker (Group 2) to identify BV-positive samples (Table 1). Owing to the strong co-existence association between *G. vaginalis, P. bivia, Megasphaera sp. type 1,* and *A. vaginalis* (Fig S10 and S23), they were tied together to serve as a marker (Group 3) to fine-tune our criteria in distinguishing between transitional BV, BV-positive, and BV-negative samples. Finally, the remaining eight species were also bundled into a marker (Group 4) to further distinguish between BV-positive, BV-negative, and transitional BV (Table 1).

**Table 1.**
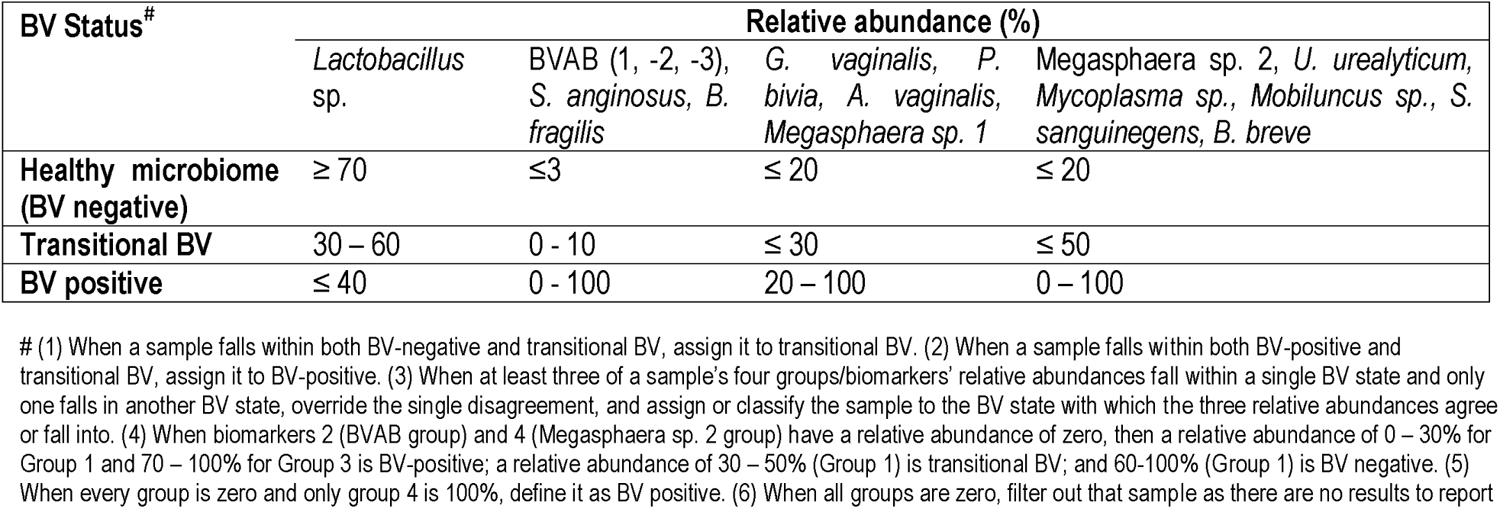
MDL-BV index designed to interpret the new 22-qPCR results and diagnose vaginal samples as BV-positive, BV-negative, or transitional BV.

The relative abundance distribution of the species within each of the four biomarker groups above were then used to select cut-off ranges for each group, incorporating a relative abundance-based criteria into the species-based criteria above. This resulted in a four-marker criterion, called the *MDL-BV index*, for diagnosing BV. We trained and tested this two-tier diagnostic *MDL-BV index* on our samples using machine-learning codes (Decision Trees and Random Forests) in Python, adjusting the ranges of each (species group) biomarker’s relative abundance until an optimal range per (species group) biomarker was found. The results of the *MDL-BV index’s* classifications were manually verified to ensure its veracity.

Observations made during the training and testing process made us include the following four instructions into the *MDL-BV index* code to enable categorization of all types of vaginal samples into their respective BV states: 1. When a sample falls within both BV-negative and transitional BV, assign it to transitional BV; 2. When a sample falls within both BV-positive and transitional BV, assign it to BV-positive; 3. When at least three of a sample’s four groups/biomarkers’ relative abundances fall within a single BV state and only one falls in another BV state, override the single disagreement, and assign or classify the sample to the BV state with which the three relative abundances agree; 4. When biomarkers 2 and 4 have a relative abundance of zero, then a relative abundance of 0 – 30% for Group 1 and 70 – 100% for Group 3 is BV-positive; a relative abundance of 30 – 50% (Group 1) is transitional BV and 60 – 100% (Group 1) is BV negative; 5. When every group is zero and only group 4 is 100%, define it as BV positive; 6. When all groups are zero, filter out that sample as there are no results to report (Table 1).

The final *MDL-BV index* was then tested on the data used in this study and 490 samples were classified as BV positive, 335 samples were classified as BV negative, and 151 samples were classified as Transitional BV. A manual verification of these classifications found them to be accurate, based on the MDL-BV index ranges. The relative abundance of each of the four biomarkers/group and the final BV status classification based on these relative abundances, produced by the Python code, is shown in Table S3 (Table 1).

### 3.6 Demographics and BV

The effect of age, pregnancy status, and race on BV status was analyzed using a correlation heatmap (Fig. S24) and Box and Whisker distribution plots (Figure 8). Notably, almost all BV-negative cases were found within the White population with virtually none being found among the other races. Transitional BV was found only among Whites and Blacks, with BV-positive cases being widely distributed among the Black population aged 28 – 40; most White women who had BV were aged between 25 and 35. The age group of the BV-positive and transitional BV population were almost the same among Blacks but very different among Whites. The ‘Other” race (including Latinos/Hispanics) also had substantial BV, with their ages falling between a tight window of 28 to 32 years. Although the number of Asian and American Indian samples were relatively few, they were all BV-positive and fell within the median age range.

**Figure 8.**
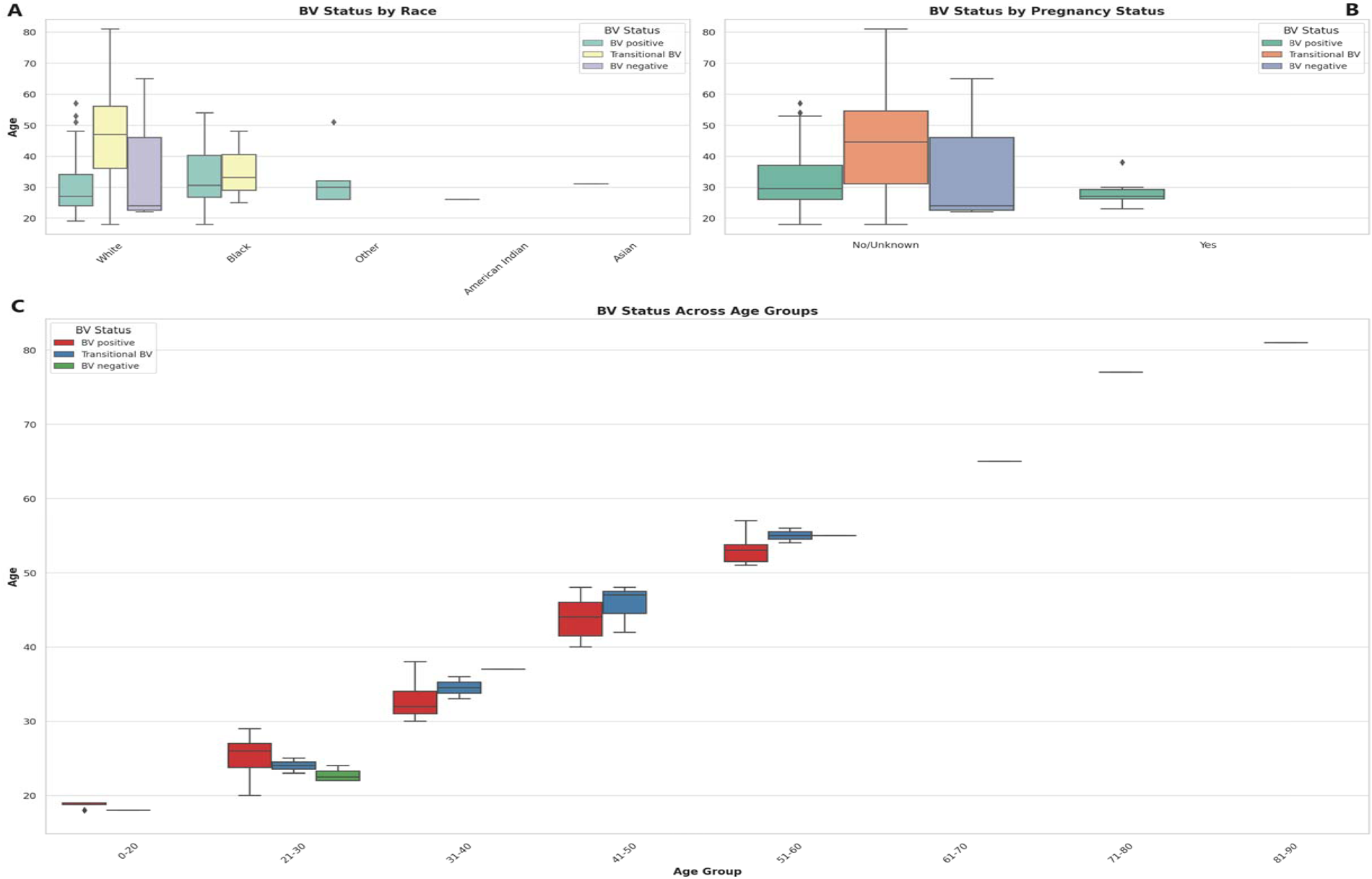
Box and Whisker plots showing the distribution of BV-positive, Transitional BV, and BV-negative samples across different races, ages, and pregnancy status. **A** shows the distribution of BV status across different races. **B** illustrates the distribution of BV status among pregnant and non-pregnant women. **C** visualizes the distribution of BV status across different age groups. The horizontal axes show the ages for the different classifications and distributions. The box colors are not the same for all the three plots and their respective keys in the upper right or left corners designate the correct interpretation of each color. The Whisker show the upper and lower quartiles while the boxes show the 25^th^ and 75^th^ percentile range (50% of the population) while the stars show outliers. Chi-square and Kruskal-Wallis tests respectively showed that there were no significant effect/association between race, pregnancy status, and age groups with BV.

It is notable that most women who were pregnant were also BV-positive, with no transitional and BV-negative status being found among them. Further, the age differences between BV-positive pregnant women (24 – 30 years) and non-pregnant women (28 – 38 years) were evidently wide. The Box plot in 8C shows a similar occurrence of BV among women aged 21 – 60, with very little incidence being found among women aged above 60 and below 20. Notably, transitional BV cases were common among women aged between 41 and 50 than among the other age groups. BV-negative cases were mostly found among women aged between 21 – 30 years.

Chi-square tests indicated that there was no statistically significant association between race and BV status (*P* = 0.217), and between pregnancy status and BV status (*P* = 0.527) at significance levels (*P* < 0.05). The Kruskal-Wallis test, which was used to assess whether there are statistically significant differences in age distributions between the BV positive and BV negative groups, found no statistically significant differences in the age distributions between the BV positive and BV negative groups (*P* = 0.820) at significance levels (*P* < 0.05). Therefore, age by itself, may not be a distinguishing factor between these two groups within the dataset analyzed.

## 4. Discussion

Unlike other infections with single etiological agents, BV is a polymicrobial infection with no clear consensus on the specific bacterial species responsible for the altered vaginal microflora state [20,23]. The exact contributions of BV-associated microbes to the pathogenesis of BV, which may be relevant for accurate diagnosis and therapeutics, remain unresolved. Nevertheless, the occurrence of normal, transitional, and abnormal vaginal microflora states is dependent on the composition and interactions of the different *Lactobacillus* and anaerobic species present [19,26–28]. To address this gap between the vaginal microbiome dynamics and the current BV diagnostic tests’ limitations, a new qRT-PCR test that detects and measures the gDNA concentrations of 22 species found in all the various conditions of the vaginal microbiome was designed.

With such a broad spectrum of bacterial species, the resolution of the test is severally enhanced to efficiently distinguish between normal, abnormal, and transitional flora. The test was able to identify all plasmid standards with 95 – 100% sensitivity and specificity with a short turnaround time of 8 hours (from sample reception). In studies where results from PCR or qPCR studies targeting 2 – 13 species have been compared with Amsel’s criteria or Nugent’s score, the results have been highly sensitive and specific [26,29–32]. We are therefore confident that subsequent studies with our test, using Nugent score or Amsel’s criteria-diagnosed samples, will equally yield similar if not better results with its larger bacteria target spectrum. Although this study is limited by the inability to compare the current results with clinical presentation data, its sensitivity and specificity in detecting the controls used confirm its efficiency.

In fact, the test’s diagnostic capability is further appreciated when the make-up and sample size of our vaginal specimens are considered: 946 vaginal samples from a wide-spectrum of ages (18– 83), races (White, Black, Asian, Indian, and “Other”, not to mention the races of most women who failed to state their race), and pregnancy status. The Bray-Curtis dissimilarity matrix and the principal component analysis (PCoA) showed how dissimilar the samples were as few of the samples clustered together (Fig. 6). The sample differences were further highlighted by the species richness (α-diversity) of the samples in which the number of species per sample ranged from 2 – 12, with 50% of the samples having 2 – 6 species per sample. Instructively, the Shannon index further clarified this observation by showing that the species diversity per sample was low, with most samples having an index between 0.8 and 1.8 and the most diverse sample having an index of 3.2 (out of 3.5).

Thus, although the samples were dissimilar in terms of composition, their diversity were relatively low, which could be characteristic of a healthy vaginal microbiome if the relative abundance of *Lactobacillus sp.* is high. [22] However, it is worth noting that the Shannon diversity is affected by the number of species sampled and the vaginal microbiome is naturally not as diverse as the gut microbiome [5,24,33]. Hence, although the 22 species used are more than any PCR test, it may not represent all the species in the vaginal flora. For instance, the alpha diversity of the samples was calculated through a direct count of the number of species observed in each sample without using any estimators or indices. Hence, it does not account for undetected species or attempt to estimate the total species richness in the samples, which methods like the Chao1 index aim to do. Notwithstanding, the representative nature of the samples used in this study is self-evident.

To our knowledge, no qPCR assay has the same broad-spectrum species target, making this assay an important innovation and the first to do so. Whereas other assays have used only two or up to 13 species [32], not all of the species were quantitatively detected as was done in this assay [28,30,32,34]. This is despite the calls for increasing the bacterial spectrum for molecular BV diagnosis, focusing on *Megasphaera, G. vaginalis, F. vaginae*, and other anaerobes[26,30,34]. This study answers that call.

By both detecting and quantifying the 22 species in vaginal samples, we were able to obtain a clearer picture of the vaginal microbiome’s composition and make a better diagnosis of its condition; a feat only achievable with whole-genome metagenomics [35–37]. For instance, although BVAB-1, -2, and -3 were detected together in the same study [38], BVAB-2 has been commonly tested as a marker of BV positivity without BVAB-1 and -3. Yet, this study showed that BVAB-2 is the least prevalent/abundant among these three species, with BVAB-1, followed by BVAB-2, being the most common. A Brazilian study using 223 BV-positive samples also reported the same findings regarding the relative abundance of BVAB-1, BVAB-2, and BVAB-3, affirming our observations [32]. A positive association between BV and high-risk Human Papilloma Virus (HPV) genotypes has been reported, owing to the occurrence of BVAB 1 and 3, and other BV-associated-bacteria in women co-infected with HIV and HPV [39]. The same study also showed that the presence of BVAB 1 and 3 had an elevated likelihood of increasing the severity of cervical neoplasia in this population [39]. Nevertheless, these species are not as closely related as initially thought as a recent phylogenetic analysis identified BVAB 1, 2, and 3 to be *Clostridiales genomosp*., *Oscillospiraceae bacterium* strain CHIC02, and *Mageeibacillus indolicus*, respectively [37].

Additionally, the relative abundance of the various species identified by this test mirrors what has been reported [2,25,32,40,41], with *Lactobacillus sp.* (except *L. acidophilus*), *G. vaginalis, A. (F.) vaginae, P. bivia,* and *Megasphaera sp. type 1* being the most dominant and widely distributed (Fig. 2, 4, S1-S22). The absence of *M. genitalium* (which was found in low abundance in other studies using BV-positive samples [32]), and lower abundance and prevalence of *L. acidophilus, M. hominis, U. urealyticum, Megasphaera* sp. type 2*, Mobiluncus sp., S. sanguinegens, B. breve,*and *S. anginosus* are also consistent with other findings [3,27,42,43], further crediting the diagnostic efficiency of this test.

It was observed from the correlation, and co-existence analysis that the five *Lactobacillus sp.* mostly co-existed together while *G. vaginalis, A. (F.) vaginae, P. bivia,* and *Megasphaera sp. type 1* also co-existed in the same samples (Fig. 4, 7, S1-S23). The Chi-square and T-tests also largely agreed with the significant association between the *Lactobacillus sp.* and the non-*Lactobacillus* species, with few exceptions (Fig. 7). This was used to form two separate groups of species-based biomarkers. Furthermore, the BVABs, *S. anginosus,* and *B. fragilis,* which are known to be mainly associated with BV microbiomes and absent in normal vaginal microbiomes [32,44–47], were also teased from the remaining non-*Lactobacillus* species and grouped into another biomarker group. The remaining eight species were then also bundled together into another fourth group (Table 1). Using the relative abundance distributions of each species (Fig. 4, S1-S22), the relative abundance range for each of the four species biomarker groups was set to form the *MDL-BV index*, which was then further trained and tested on the data using machine-learning algorithms (Decision Trees and Random Forests) to diagnose BV. A relative abundance-based approach was adopted over a nominal concentration value because concentrations vary from sample to sample, and the swabbing method of sample collection among clinicians is not standardized; hence, samples collected by each swab is not quantitative.

The *MDL-BV index* has a high cut-off range for negative BV status (≥70% *Lactobacillus sp.* relative abundance) to ensure that samples diagnosed as normal were BV-negative in verity. It also errs on the side of caution by classifying samples qualifying as both BV-negative and transitional or transitional and BV-positive, as transitional BV or BV-positive respectively. The training of the model on the data further revealed certain intricacies and nuances in the distribution of the species and their relative abundances, which were used to refine it by including more details and rules. For instance, in situations where only two of the four biomarkers had a relative abundance score, the ratios were adjusted (Table 1). This strengthened the diagnostic efficiency of the *MDL-BV index*, resulting in a final classification of the samples as 491 BV-positive, 318 BV-negative, and 137 transitional BV. A manual verification of the results confirmed the veracity of the classifications (Table S3).

Instead of using just two biomarkers involving all *Lactobacillus sp.* and all non-*Lactobacillus* species [22], this index uses four biomarkers, which further refines the ability of the index to correctly distinguish between transitional BV or BV-positive microbiomes. Evidently, a correct molecular diagnosis of BV is critical to its treatment and monitoring, making this test and index very important in gynecology and obstetrics. The next stage of this research is to train the model on large datasets with Nugent scores/Amsel criteria as a means of further enhancing it and strengthening its proof of concept in BV diagnosis.

The distribution of BV among the different demographic variables viz., age, race, and pregnancy status showed important differences, albeit none of the differences were statistically significant (Fig. 8). Specifically, although the number of vaginal samples from Blacks was lower than that from Whites, there were more BV-positive samples from the former than from the latter. While most of the vaginal samples from Whites were BV-negative, those from Blacks were either BV-positive or transitional. Notably, the samples from the “Other” races (which includes Hispanics/Latinos), Asians, and Native Indians, although fewer in number, were also BV-positive, with none being negative. Furthermore, the mean age and age range for Whites with transitional BV were higher than those of all the other races, including the mean age of Whites with abnormal vaginal microbiomes (Fig. 8). Hence, although these differences were not statistically significant, they concur with other studies that show a higher prevalence of BV among Black and Hispanics than among Whites and Asians [13]. This agreement between our data and other studies further confirms the accuracy of our *MDL-BV index*.

Although BV among non-pregnant women was more prevalent than among pregnant women, BV-negative or transitional BV samples were not obtained from pregnant women (Fig. 8). This is a concerning observation as BV in pregnancy is associated with preterm labor, low birth weight, premature rupture of membranes, miscarriage, chorioamnionitis, and birth asphyxia [4,12,40]. Although the pregnancy stage of the women from whom the samples were obtained is unknown, it is known that the first trimester vaginal microbiome is similar to a BV-positive microbiome while the 2^nd^ and 3^rd^ trimesters are similar to a normal vaginal microbiome [48]. Hence, the stage of the pregnancy and vaginal symptoms should be considered in making a final therapeutic decision regarding pregnant women when using molecular-based tests. Furthermore, the difference in the number of pregnant women (n = 53) vis-a-viz the number of non-pregnant (or unknown pregnancy status) women is large and can skew the data. This is a major limitation in our data set and should be considered when analyzing the pregnancy data.

Although BV-positive samples were found in all age groups (18 – 83 years), they were mostly prevalent among women who were between 20 and 60 years, with a higher number of BV-positive and transitional BV samples being found among women between 41 – 50 years. Indeed, the median age found in this study’s population is similar to what was found in Brazil [32] and in the USA [14], with women above 21 years being more likely to suffer recurrence of BV [9,10]. Hence, the age groups found in our data concur with that of other studies, which confirms that BV is common among reproductive-age women. However, age was not significantly associated with BV occurrence in this study.

## 5. Conclusion

The new qRT-PCR assay developed by MDL is the first of its kind to quantitatively detect 22 species in the vaginal microbiome. We developed a novel classification system, the new *MDL-BV index*, which was trained on large sets of data and based on four species-based and relative abundance-based biomarkers. This multifaceted two-tier approach of using species and relative abundance provides a better diagnostic resolution for a polymicrobial infection such as BV. We are working to extend this approach to apply to other microbiome-based diagnostics to make disease diagnosis reflective of the clinical presentations. Our study is, however, limited by the absence of a clinical Nugent score or Amsel’s criteria; however, we are working to include these in the next studies that will involve the *MDL-BV index* and the 22-species qRT-PCR BV test. The clinical and socio-economic importance of this novel proprietary BV diagnostic test in obstetrics and gynecology is notable.

## Funding

This study was funded by Medical Diagnostic Laboratories, LLC (within the Genesis Global Group), Hamilton Township, New Jersey.

## Acknowledgements

The authors are grateful to the technicians at Medical Diagnostic Laboratories for their direct and indirect assistance. The material and financial resources provided by Medical Diagnostic Laboratories towards this project are warmly acknowledged and deeply appreciated. We are also grateful to Annette Daughtry for assisting with the review of the initial draft.

## Transparency declaration

The authors are scientists working at Medical Diagnostic Laboratories, LLC.

## Ethics statement

This study was conducted in accordance with the Declaration of Helsinki. This study followed the Strengthening the Reporting of Observational Studies in Epidemiology (STROBE) reporting guideline for cohort studies.

---

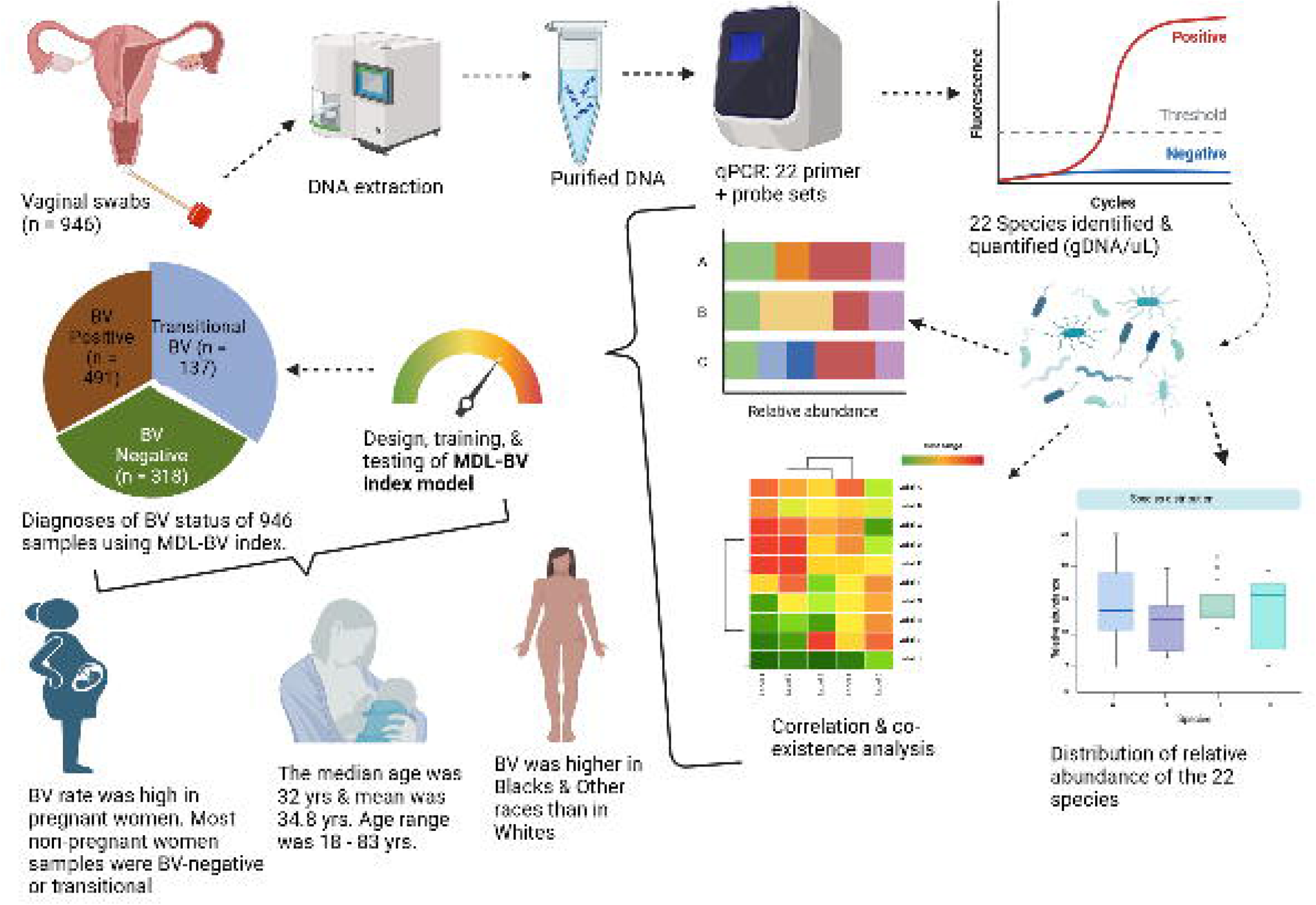

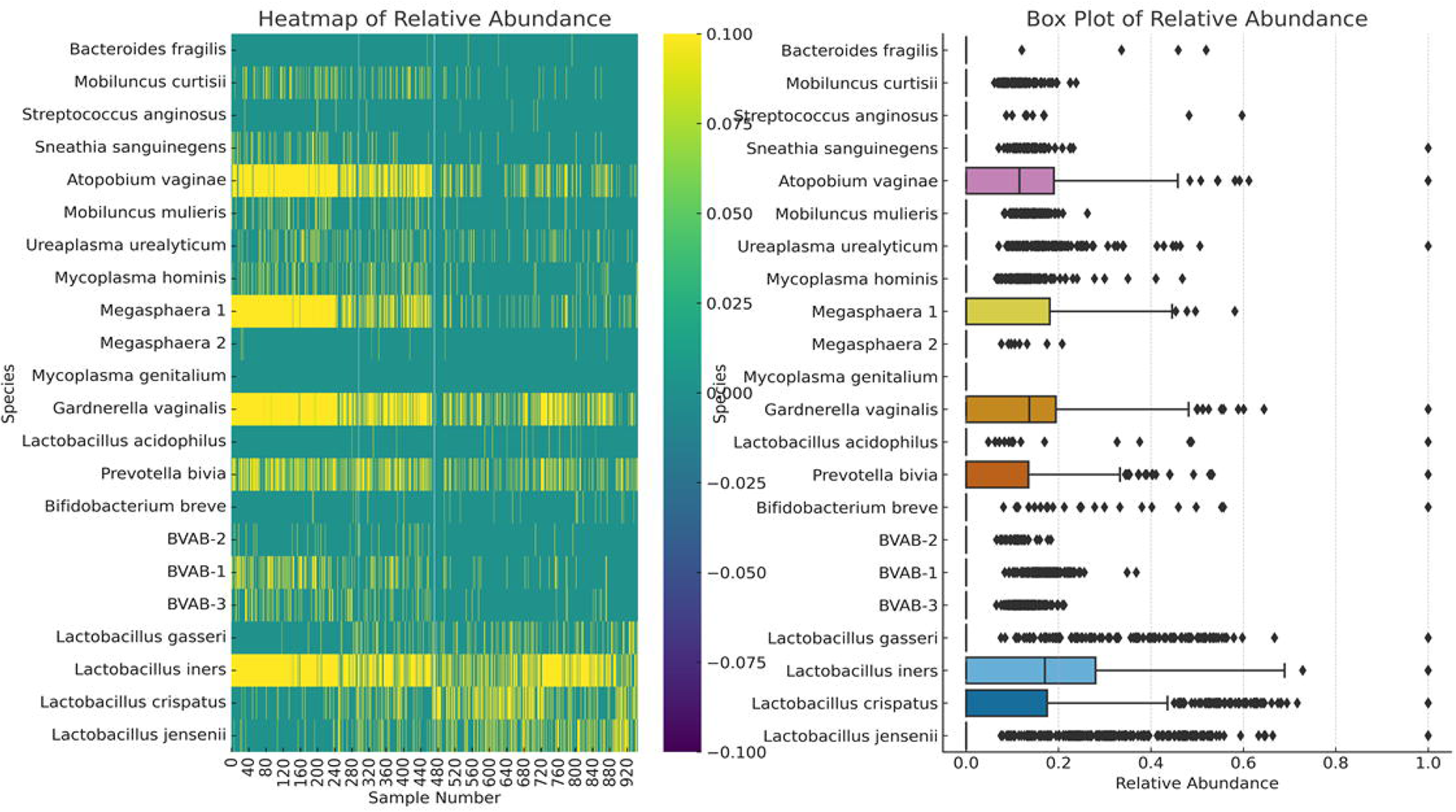

