## Supplemental figures S1-S24 for "Molecular Characterization of Vaginal Microbiota Using a New 22-Species qRT-PCR Test to Achieve a Relative-abundance and Species-based Diagnosis of Bacterial Vaginosis"

### Slide 1
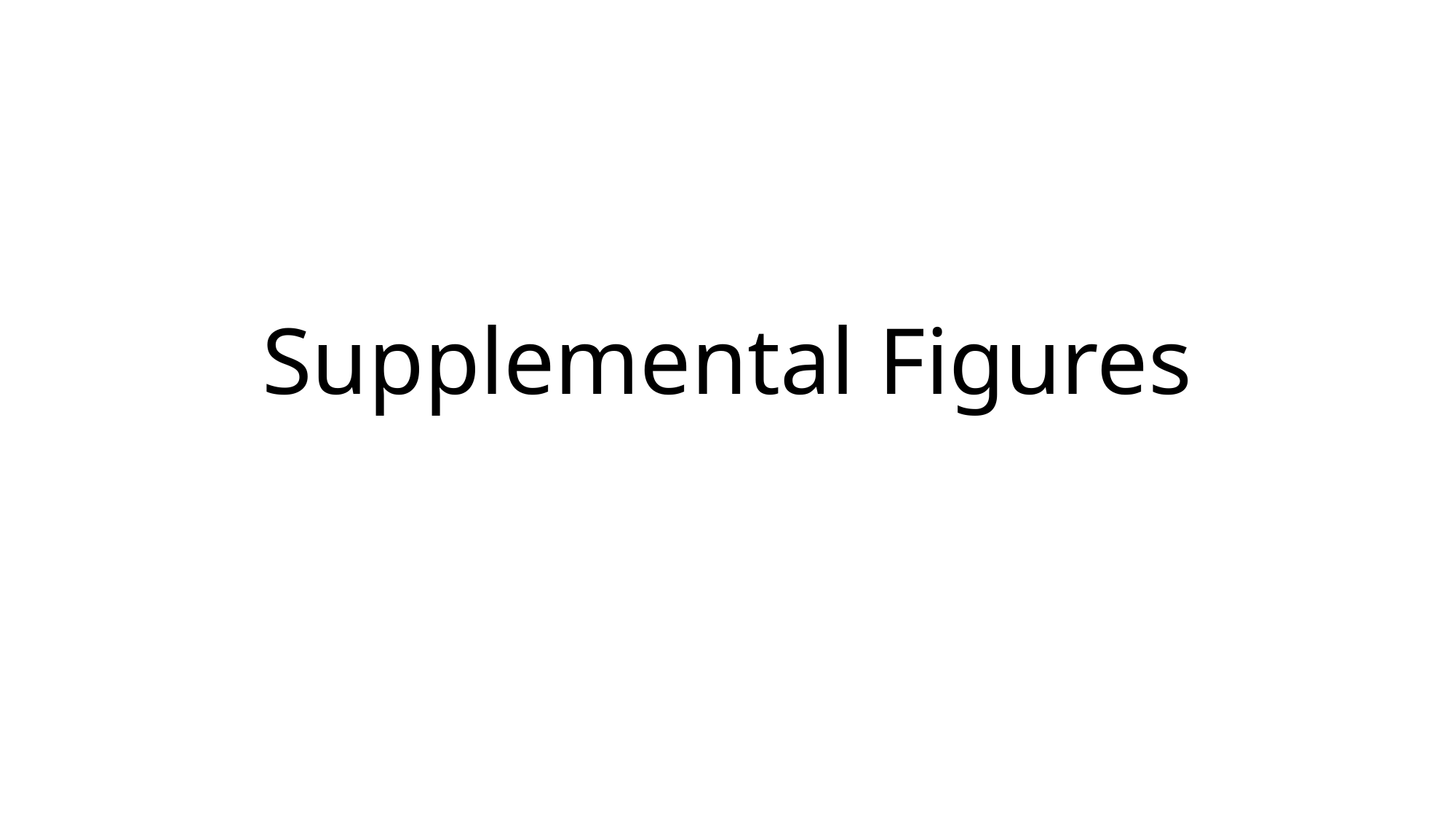

#### Supplemental Figures

### Slide 2
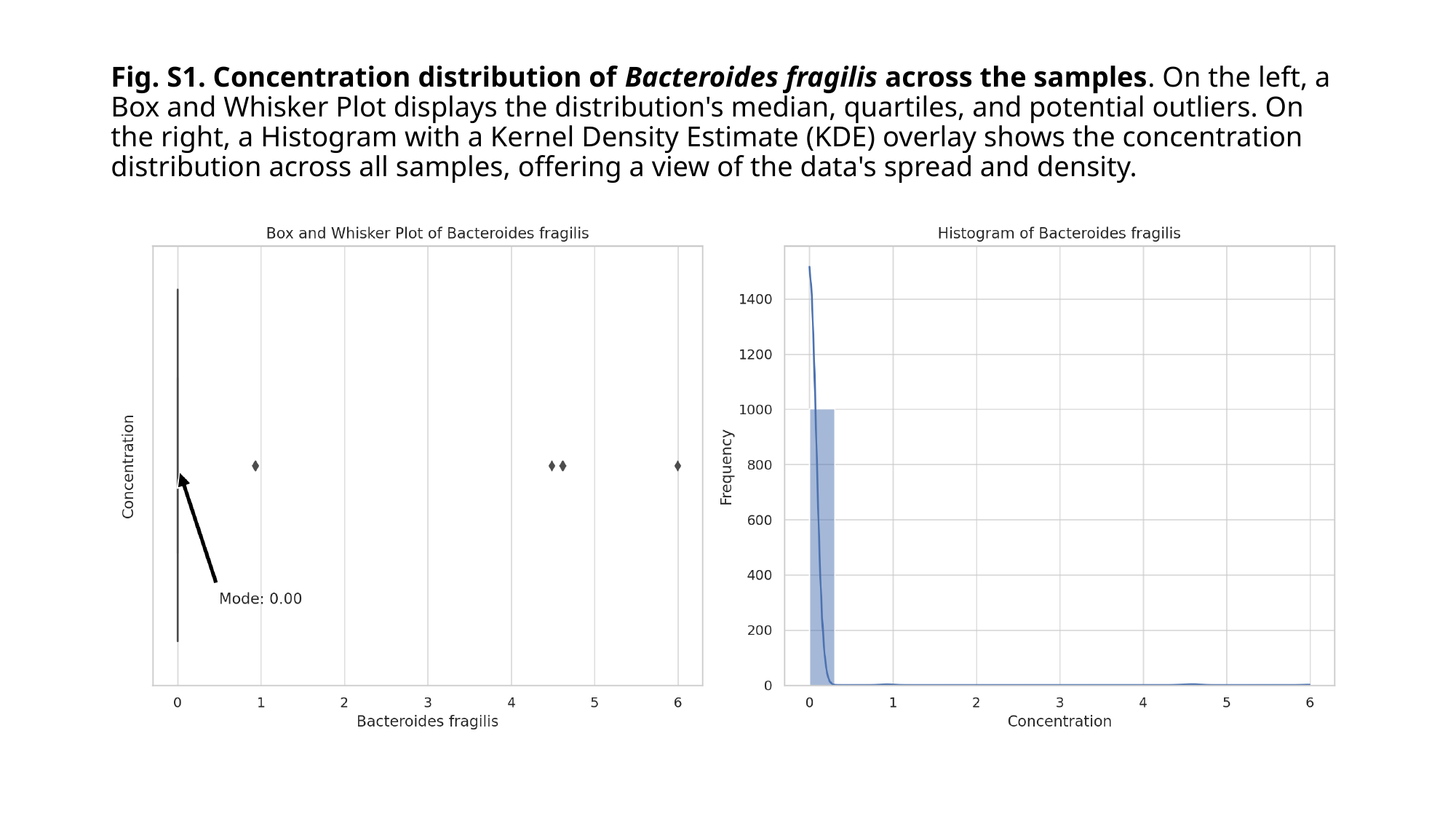

#### Fig. S1. Concentration distribution of Bacteroides fragilis across the samples. On the left, a Box and Whisker Plot displays the distribution's median, quartiles, and potential outliers. On the right, a Histogram with a Kernel Density Estimate (KDE) overlay shows the concentration distribution across all samples, offering a view of the data's spread and density.

### Slide 3
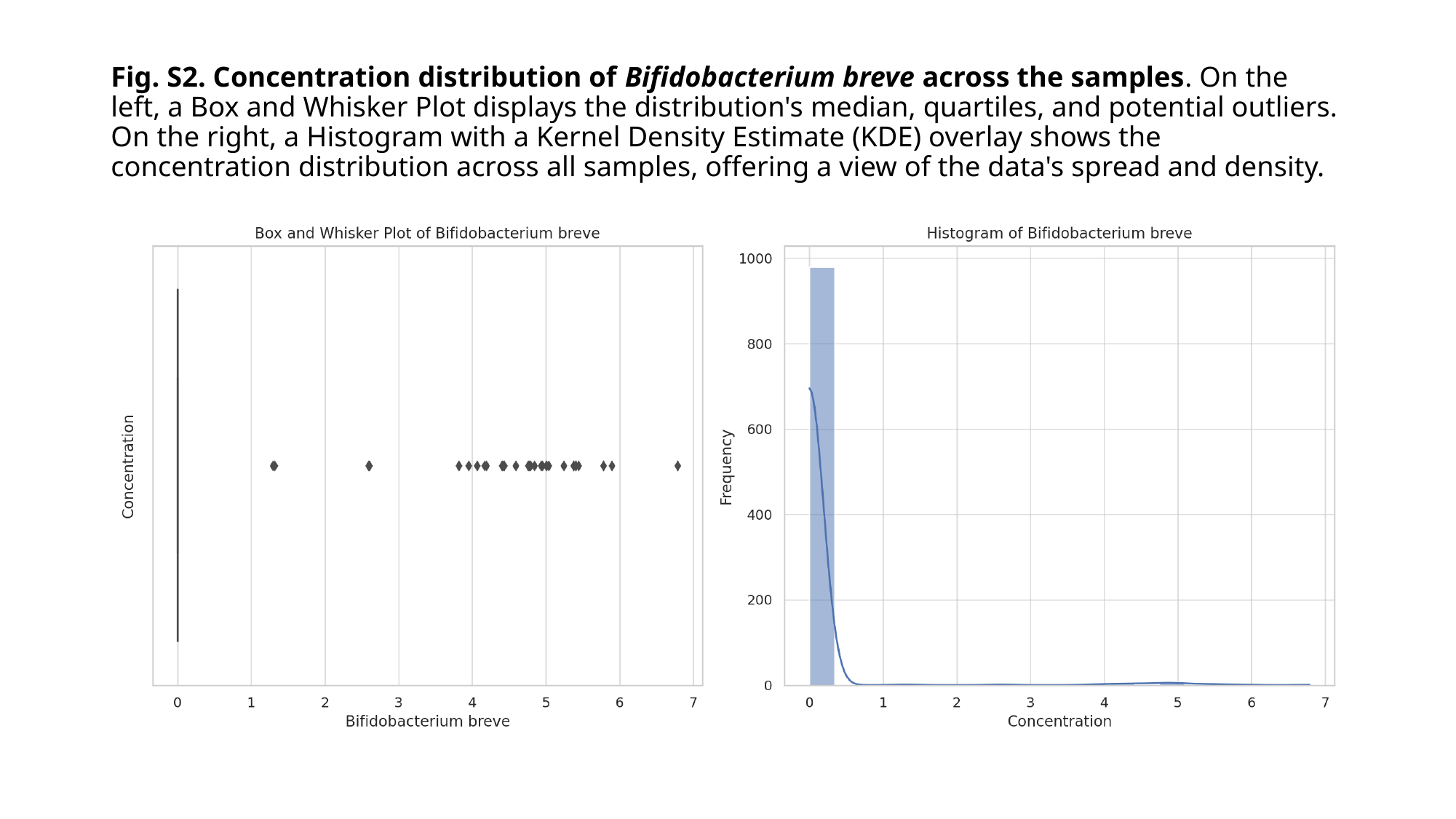

#### Fig. S2. Concentration distribution of Bifidobacterium breve across the samples. On the left, a Box and Whisker Plot displays the distribution's median, quartiles, and potential outliers. On the right, a Histogram with a Kernel Density Estimate (KDE) overlay shows the concentration distribution across all samples, offering a view of the data's spread and density.

### Slide 4
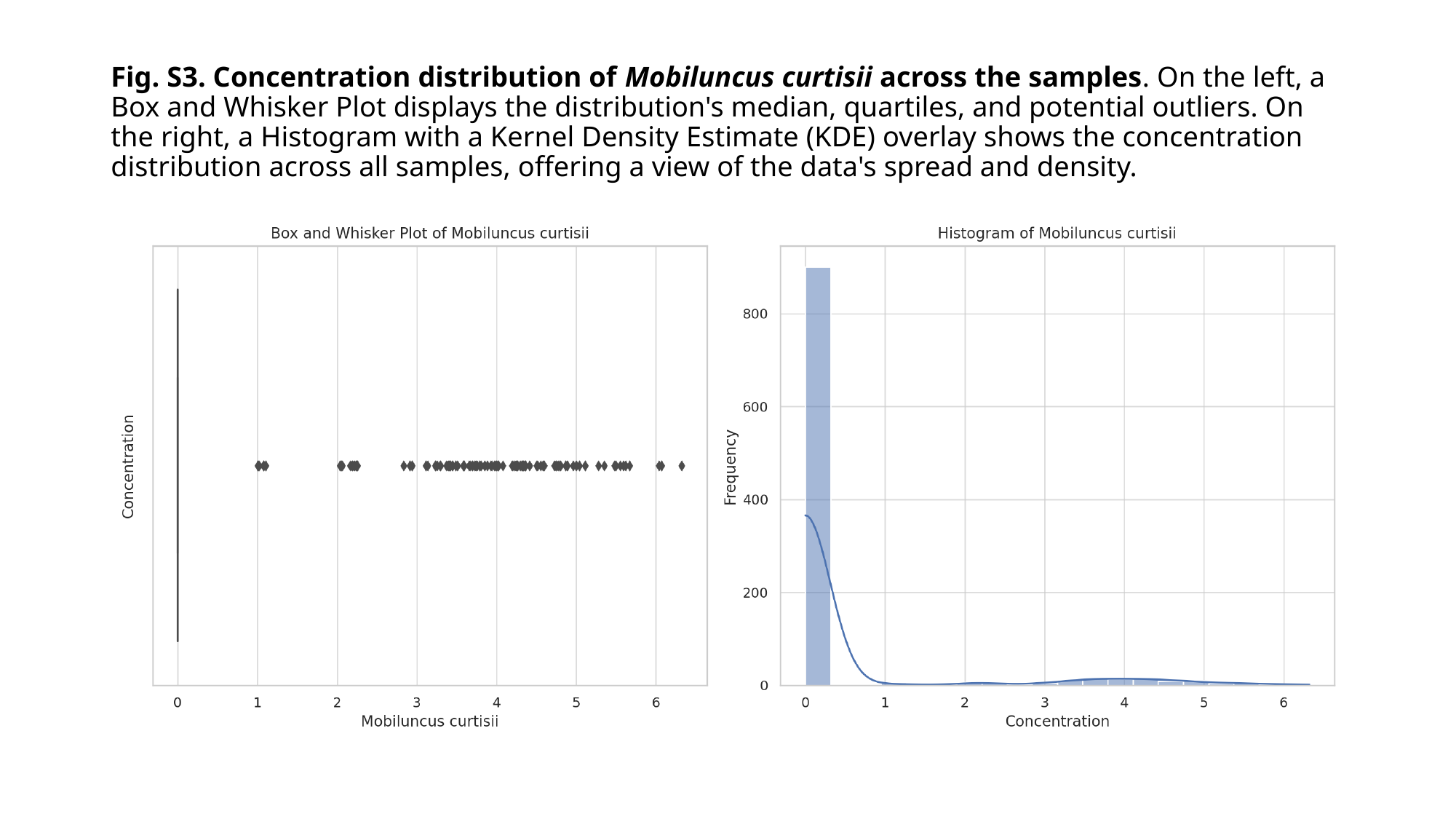

#### Fig. S3. Concentration distribution of Mobiluncus curtisii across the samples. On the left, a Box and Whisker Plot displays the distribution's median, quartiles, and potential outliers. On the right, a Histogram with a Kernel Density Estimate (KDE) overlay shows the concentration distribution across all samples, offering a view of the data's spread and density.

### Slide 5
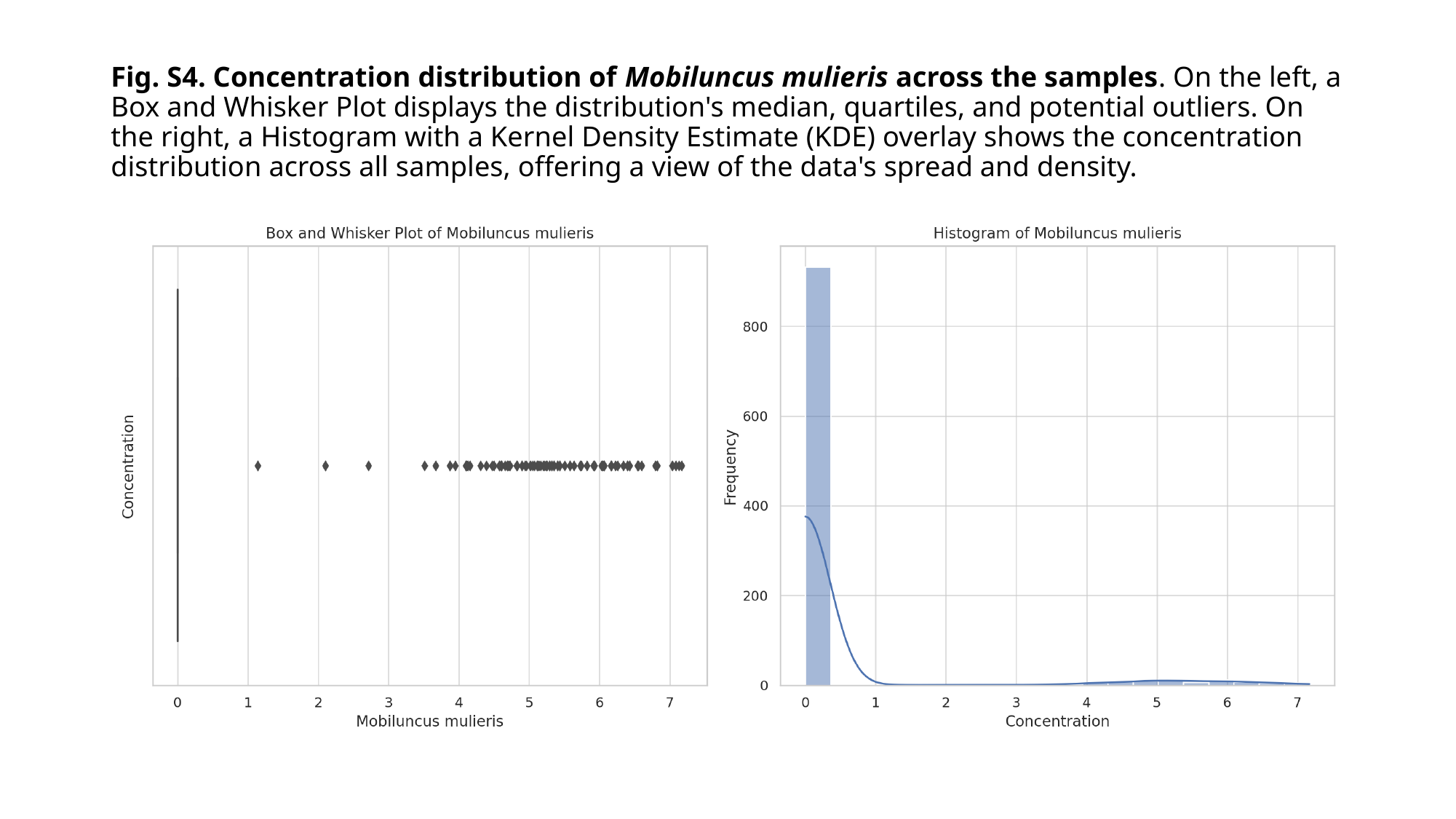

#### Fig. S4. Concentration distribution of Mobiluncus mulieris across the samples. On the left, a Box and Whisker Plot displays the distribution's median, quartiles, and potential outliers. On the right, a Histogram with a Kernel Density Estimate (KDE) overlay shows the concentration distribution across all samples, offering a view of the data's spread and density.

### Slide 6
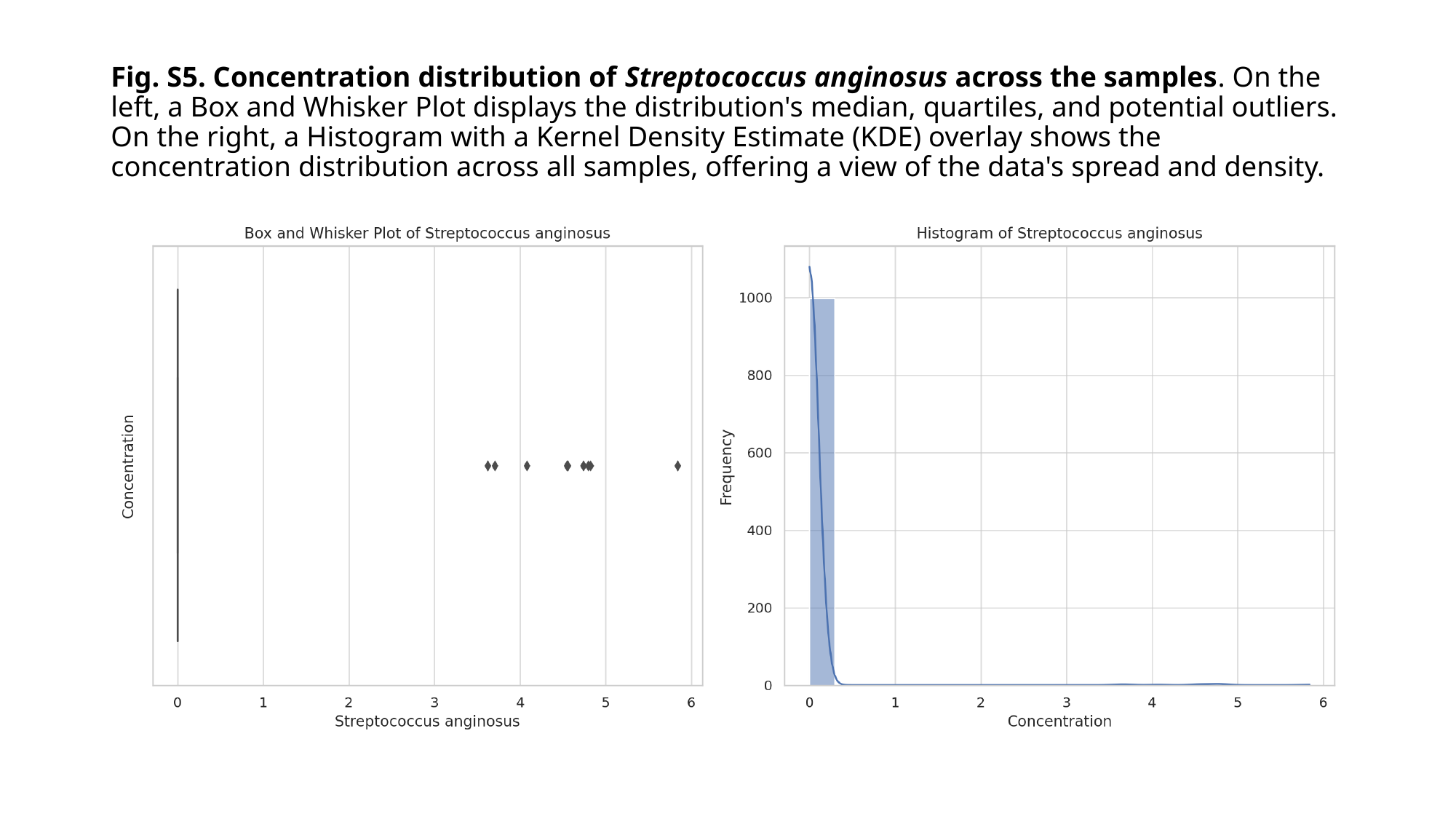

#### Fig. S5. Concentration distribution of Streptococcus anginosus across the samples. On the left, a Box and Whisker Plot displays the distribution's median, quartiles, and potential outliers. On the right, a Histogram with a Kernel Density Estimate (KDE) overlay shows the concentration distribution across all samples, offering a view of the data's spread and density.

### Slide 7
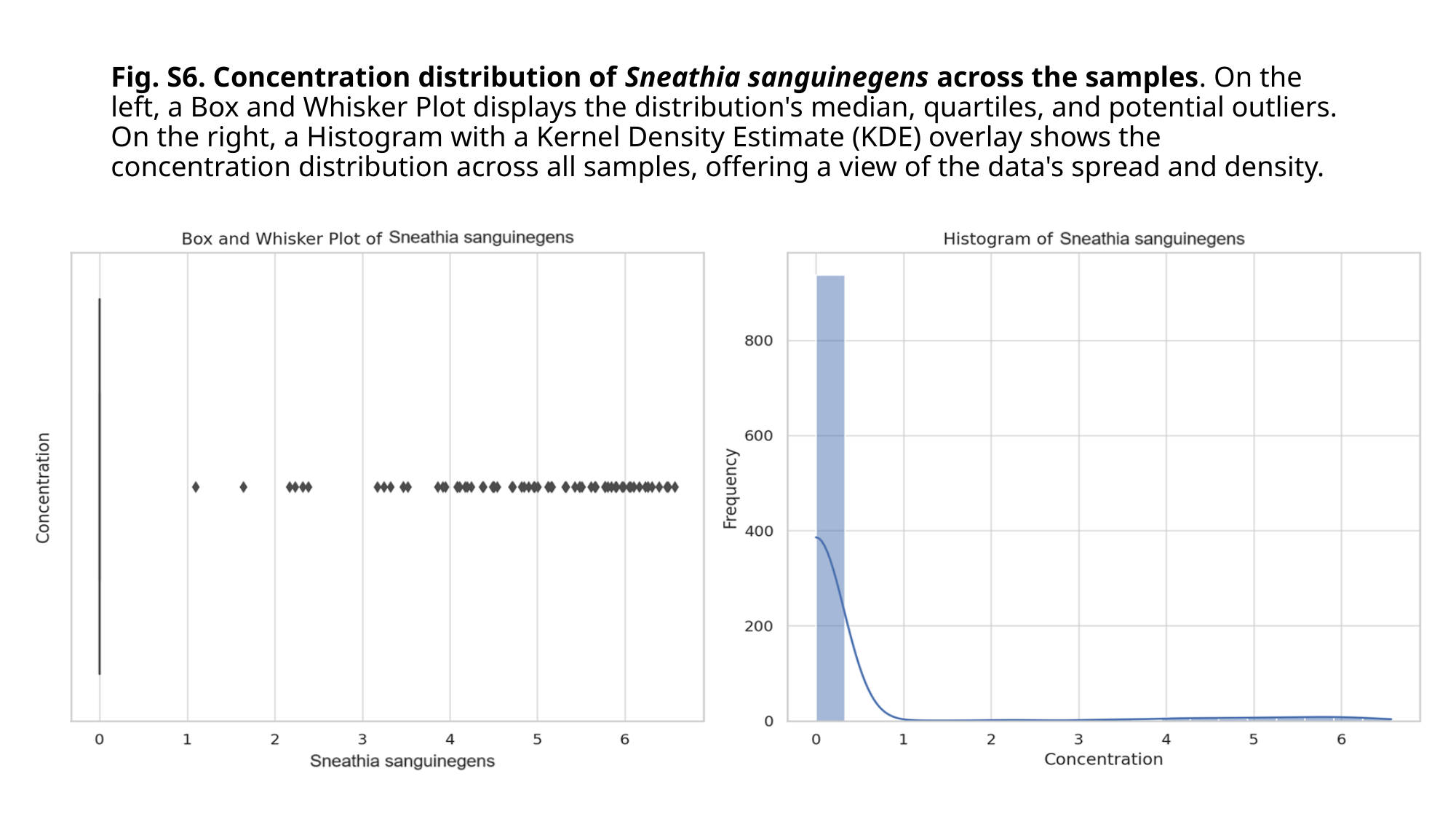

#### Fig. S6. Concentration distribution of Sneathia sanguinegens across the samples. On the left, a Box and Whisker Plot displays the distribution's median, quartiles, and potential outliers. On the right, a Histogram with a Kernel Density Estimate (KDE) overlay shows the concentration distribution across all samples, offering a view of the data's spread and density.

### Slide 8
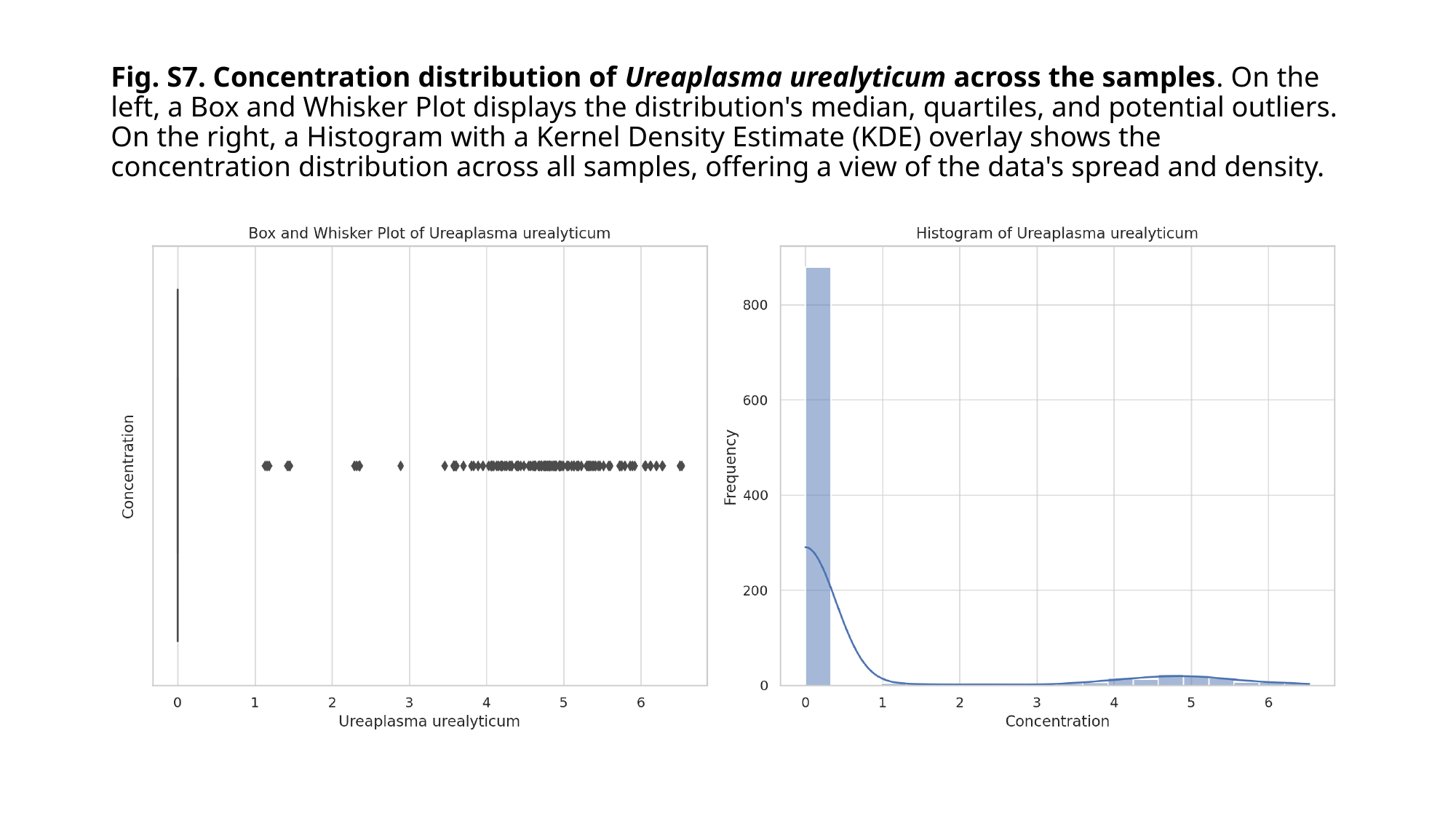

#### Fig. S7. Concentration distribution of Ureaplasma urealyticum across the samples. On the left, a Box and Whisker Plot displays the distribution's median, quartiles, and potential outliers. On the right, a Histogram with a Kernel Density Estimate (KDE) overlay shows the concentration distribution across all samples, offering a view of the data's spread and density.

### Slide 9
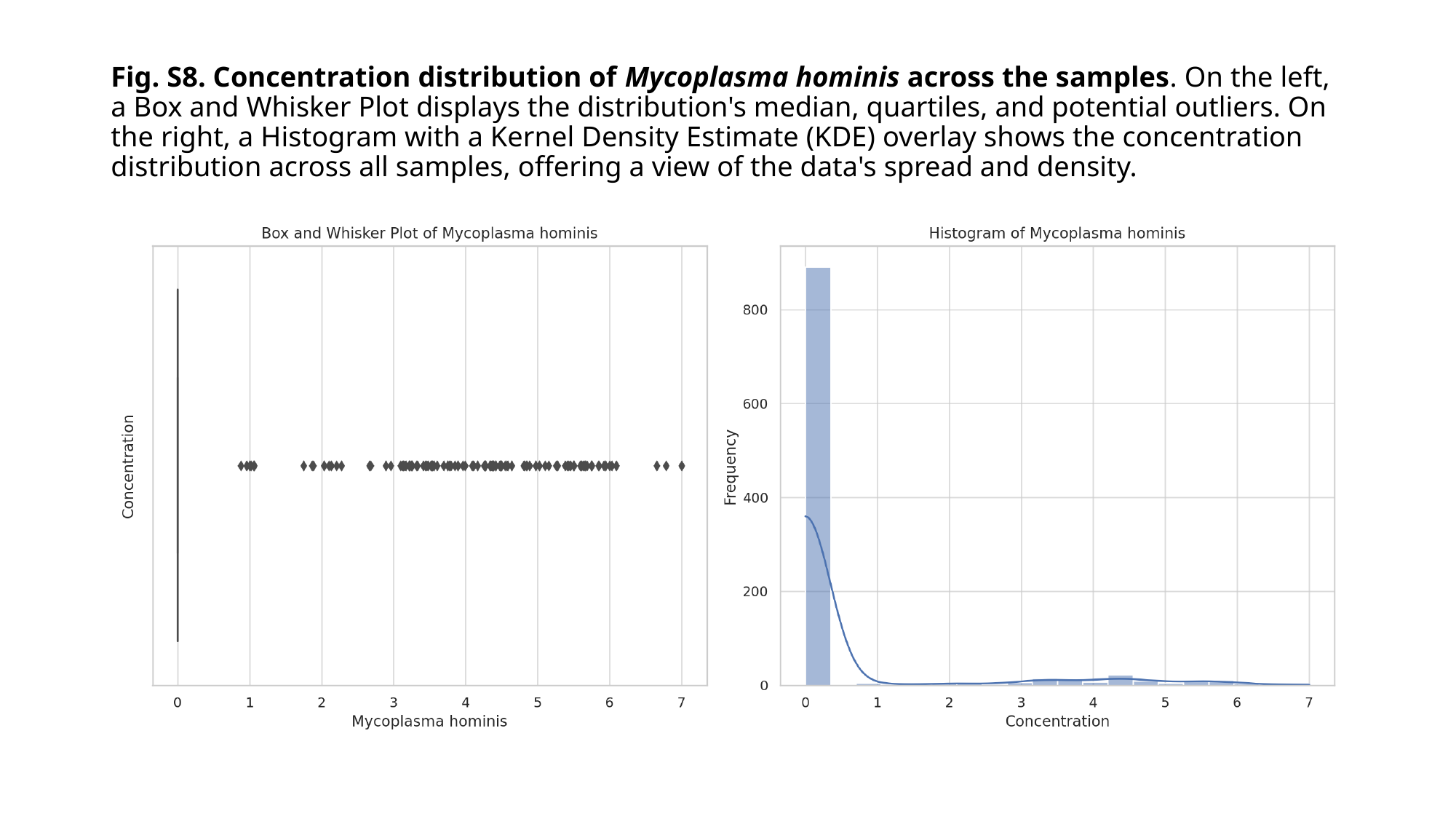

#### Fig. S8. Concentration distribution of Mycoplasma hominis across the samples. On the left, a Box and Whisker Plot displays the distribution's median, quartiles, and potential outliers. On the right, a Histogram with a Kernel Density Estimate (KDE) overlay shows the concentration distribution across all samples, offering a view of the data's spread and density.

### Slide 10
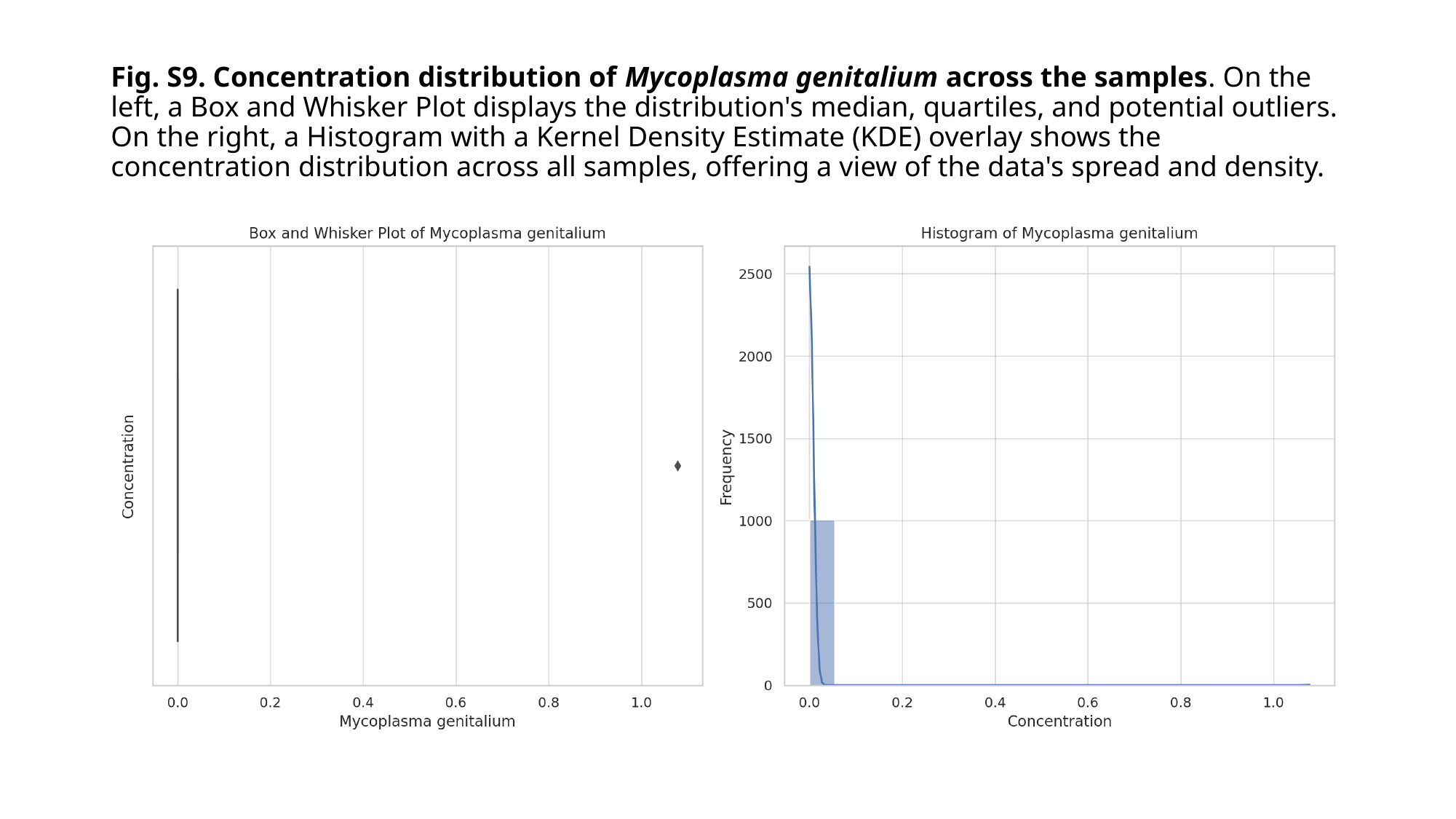

#### Fig. S9. Concentration distribution of Mycoplasma genitalium across the samples. On the left, a Box and Whisker Plot displays the distribution's median, quartiles, and potential outliers. On the right, a Histogram with a Kernel Density Estimate (KDE) overlay shows the concentration distribution across all samples, offering a view of the data's spread and density.

### Slide 11
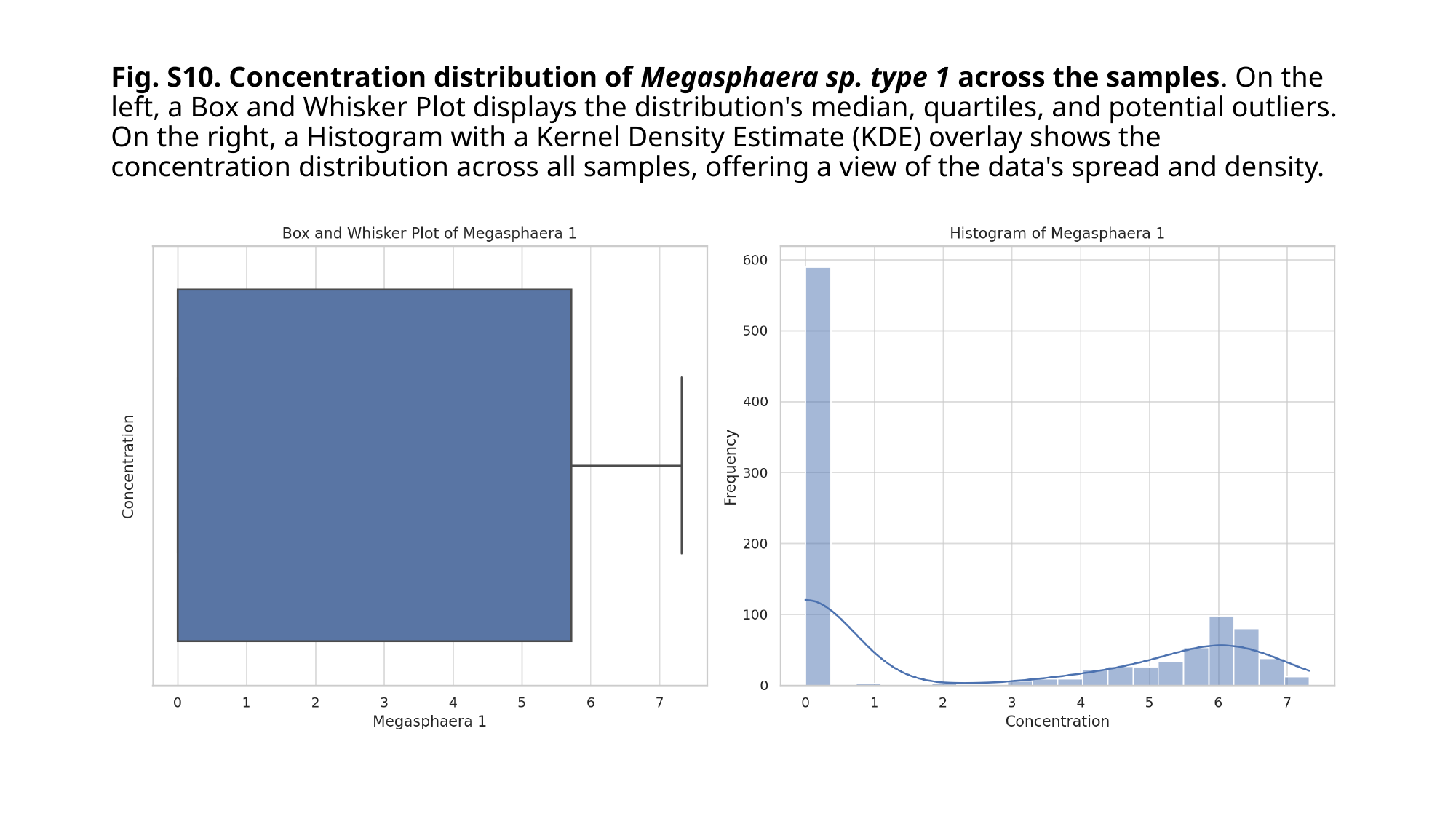

#### Fig. S10. Concentration distribution of Megasphaera sp. type 1 across the samples. On the left, a Box and Whisker Plot displays the distribution's median, quartiles, and potential outliers. On the right, a Histogram with a Kernel Density Estimate (KDE) overlay shows the concentration distribution across all samples, offering a view of the data's spread and density.

### Slide 12
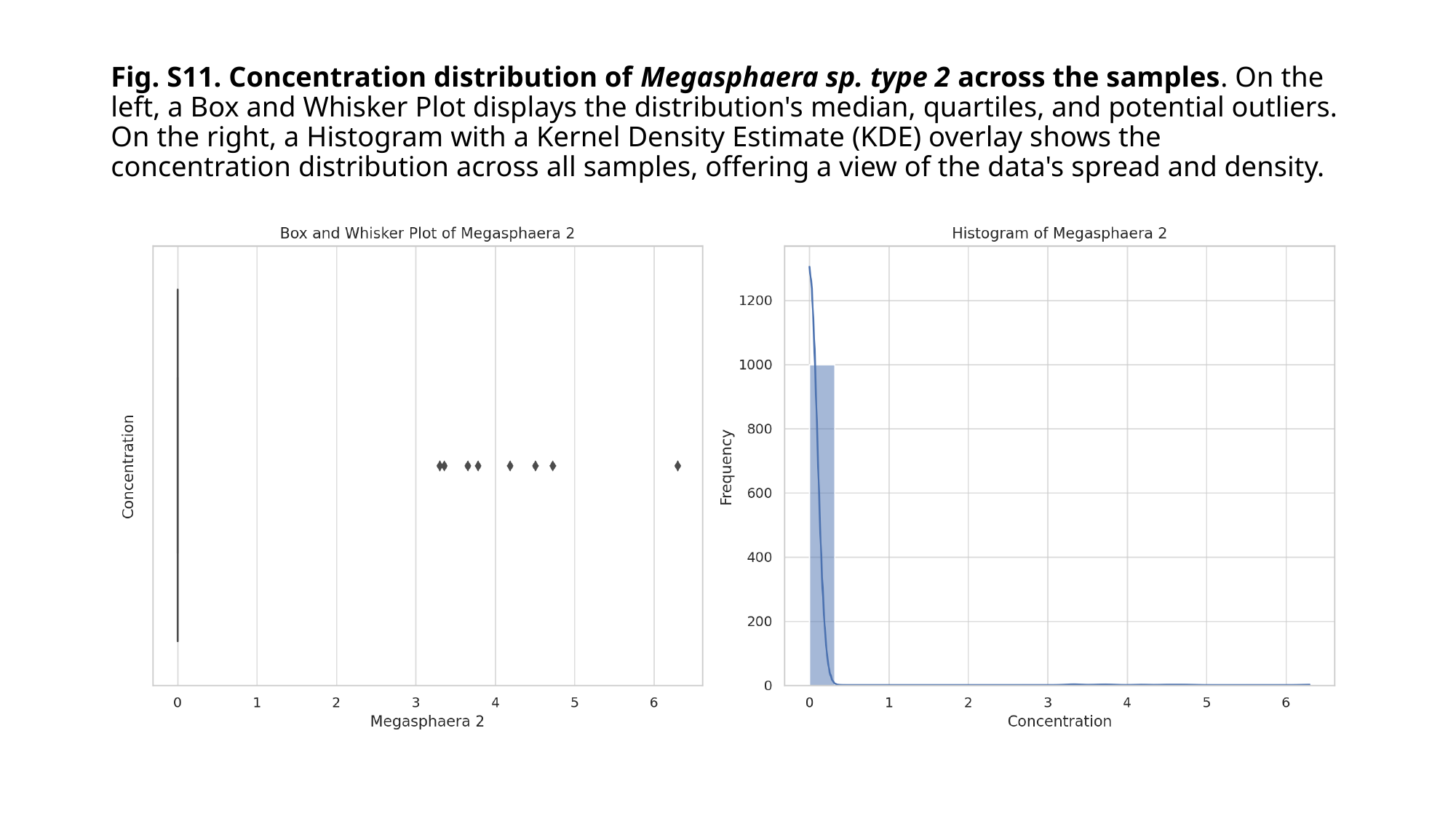

#### Fig. S11. Concentration distribution of Megasphaera sp. type 2 across the samples. On the left, a Box and Whisker Plot displays the distribution's median, quartiles, and potential outliers. On the right, a Histogram with a Kernel Density Estimate (KDE) overlay shows the concentration distribution across all samples, offering a view of the data's spread and density.

### Slide 13
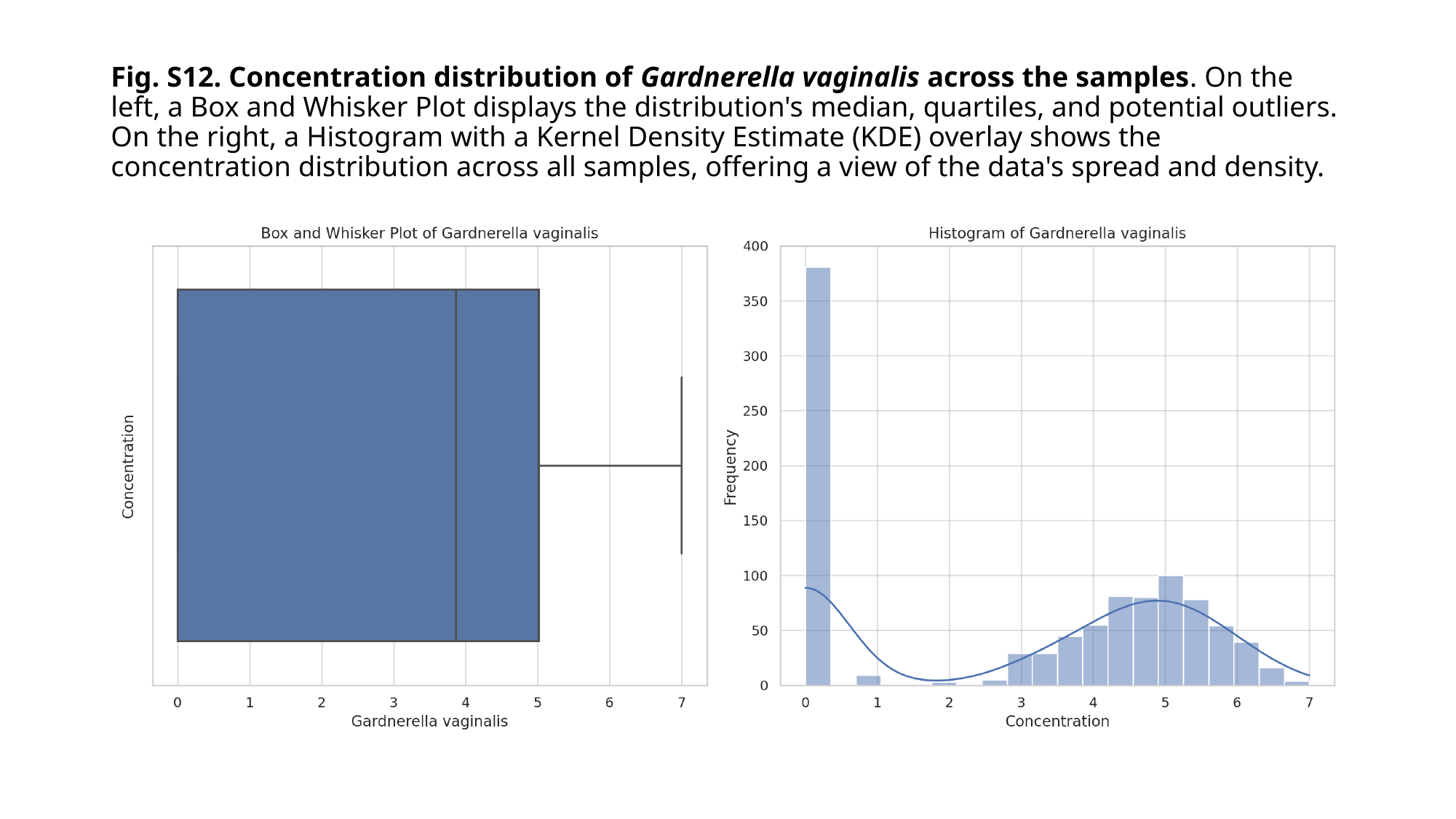

#### Fig. S12. Concentration distribution of Gardnerella vaginalis across the samples. On the left, a Box and Whisker Plot displays the distribution's median, quartiles, and potential outliers. On the right, a Histogram with a Kernel Density Estimate (KDE) overlay shows the concentration distribution across all samples, offering a view of the data's spread and density.

### Slide 14
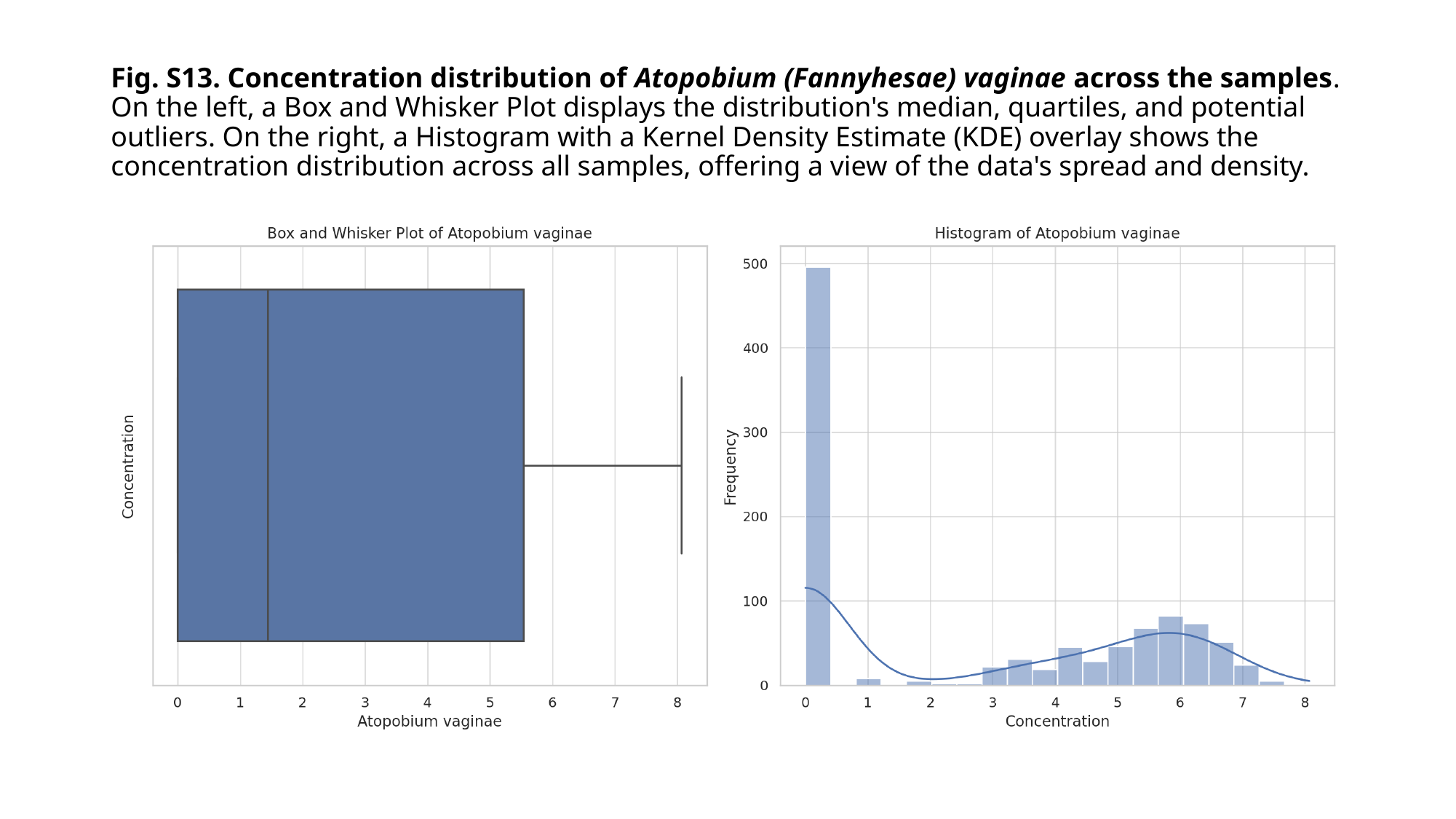

#### Fig. S13. Concentration distribution of Atopobium (Fannyhesae) vaginae across the samples. On the left, a Box and Whisker Plot displays the distribution's median, quartiles, and potential outliers. On the right, a Histogram with a Kernel Density Estimate (KDE) overlay shows the concentration distribution across all samples, offering a view of the data's spread and density.

### Slide 15
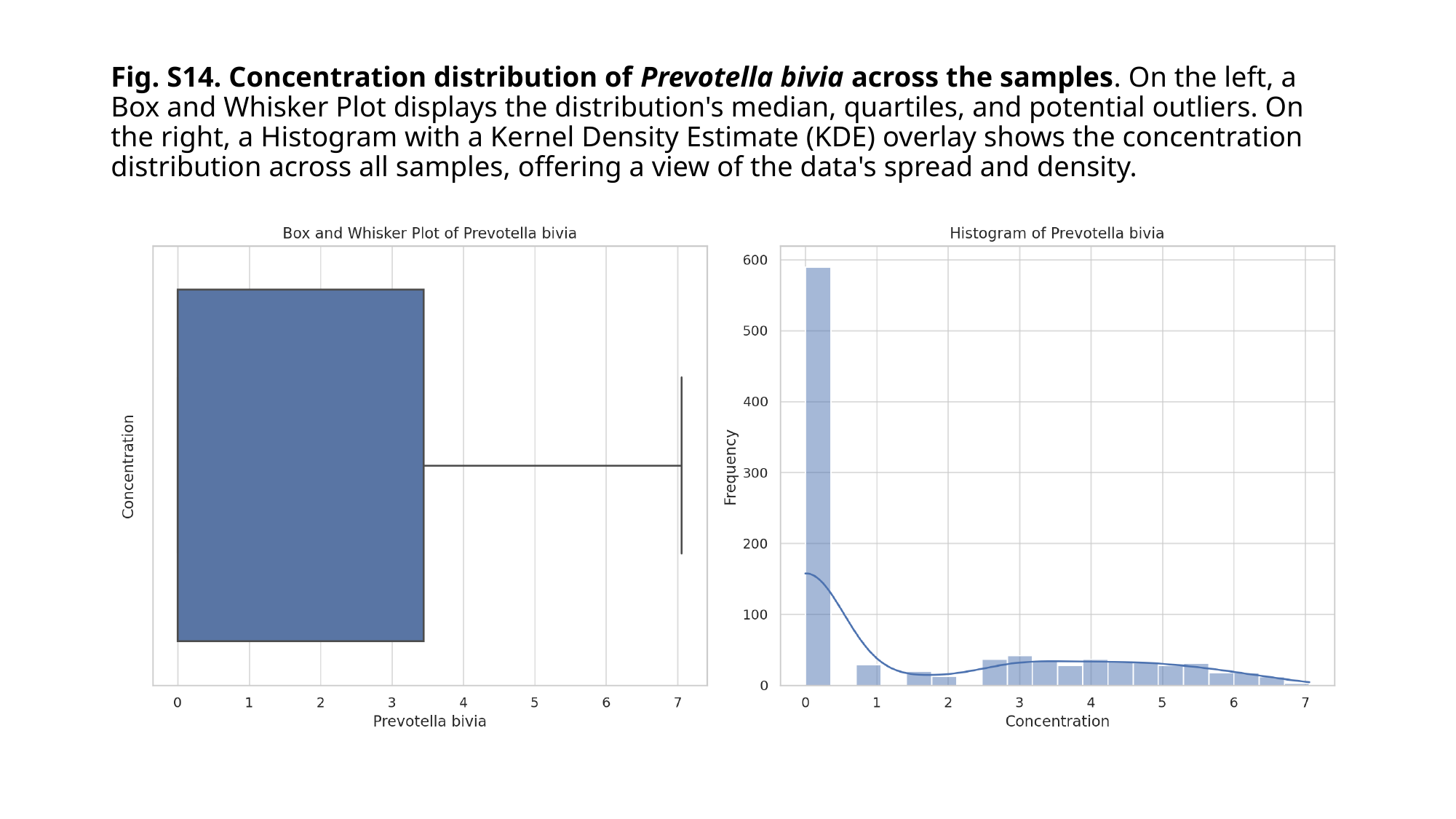

#### Fig. S14. Concentration distribution of Prevotella bivia across the samples. On the left, a Box and Whisker Plot displays the distribution's median, quartiles, and potential outliers. On the right, a Histogram with a Kernel Density Estimate (KDE) overlay shows the concentration distribution across all samples, offering a view of the data's spread and density.

### Slide 16
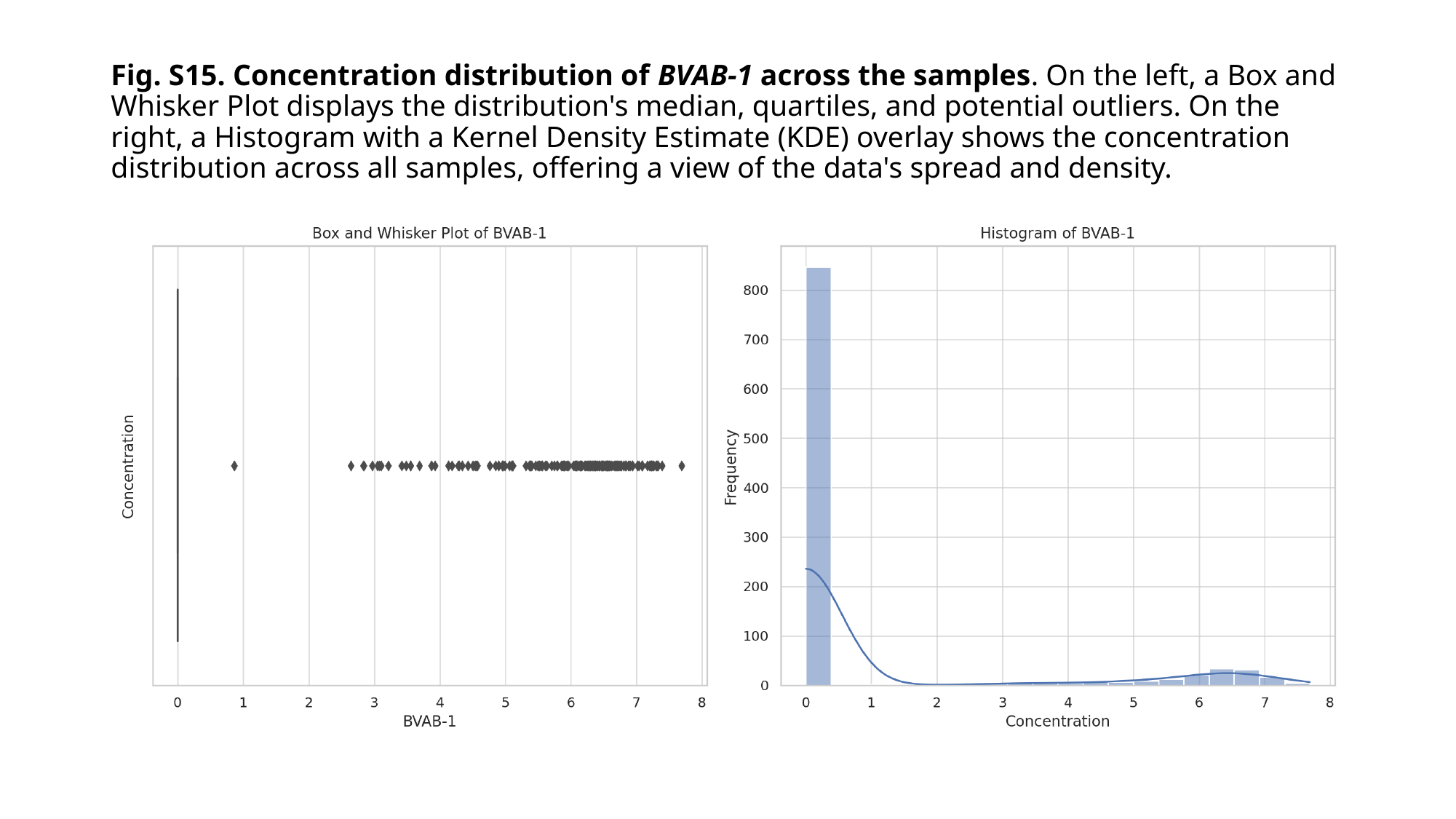

#### Fig. S15. Concentration distribution of BVAB-1 across the samples. On the left, a Box and Whisker Plot displays the distribution's median, quartiles, and potential outliers. On the right, a Histogram with a Kernel Density Estimate (KDE) overlay shows the concentration distribution across all samples, offering a view of the data's spread and density.

### Slide 17
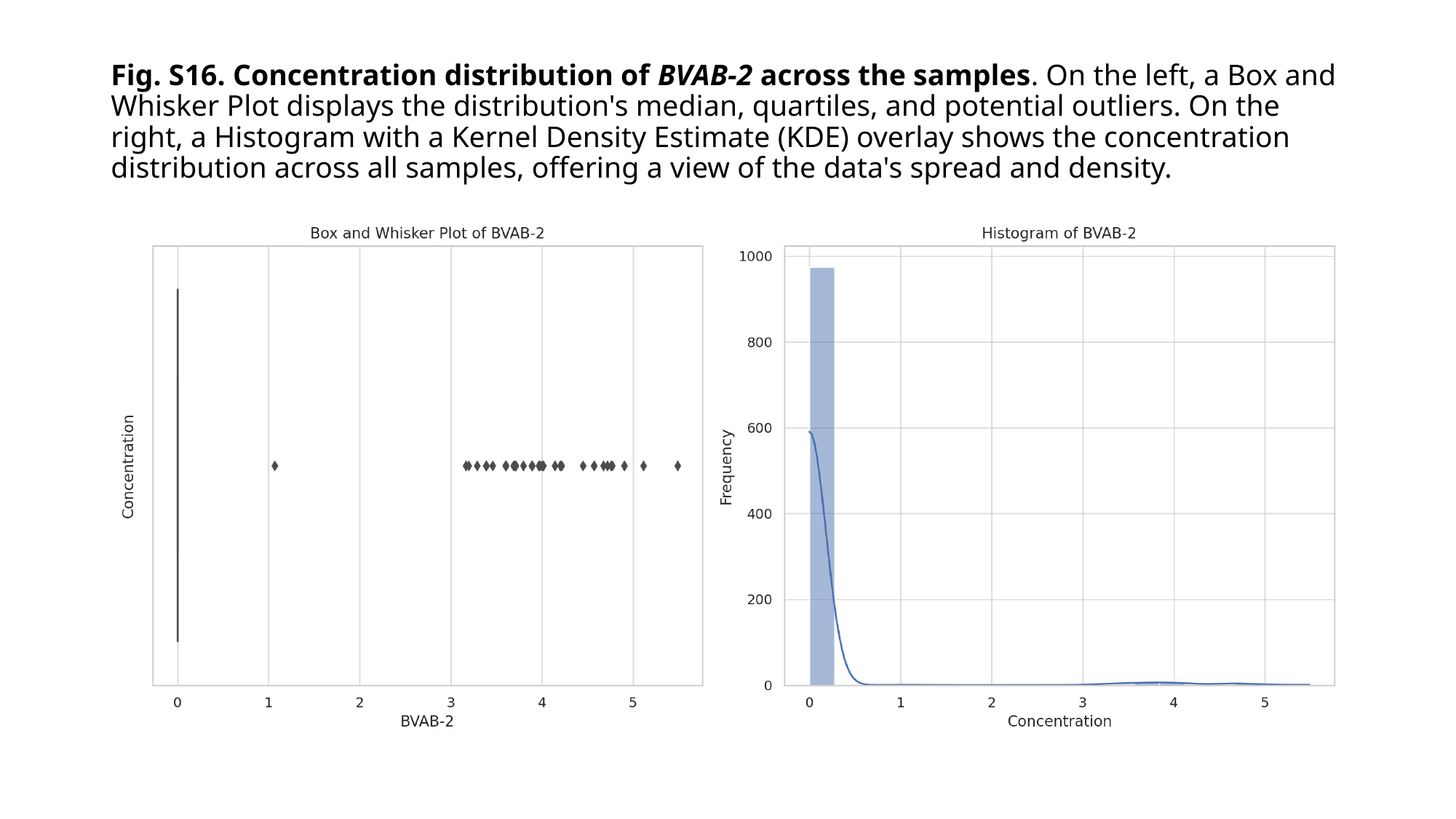

#### Fig. S16. Concentration distribution of BVAB-2 across the samples. On the left, a Box and Whisker Plot displays the distribution's median, quartiles, and potential outliers. On the right, a Histogram with a Kernel Density Estimate (KDE) overlay shows the concentration distribution across all samples, offering a view of the data's spread and density.

### Slide 18
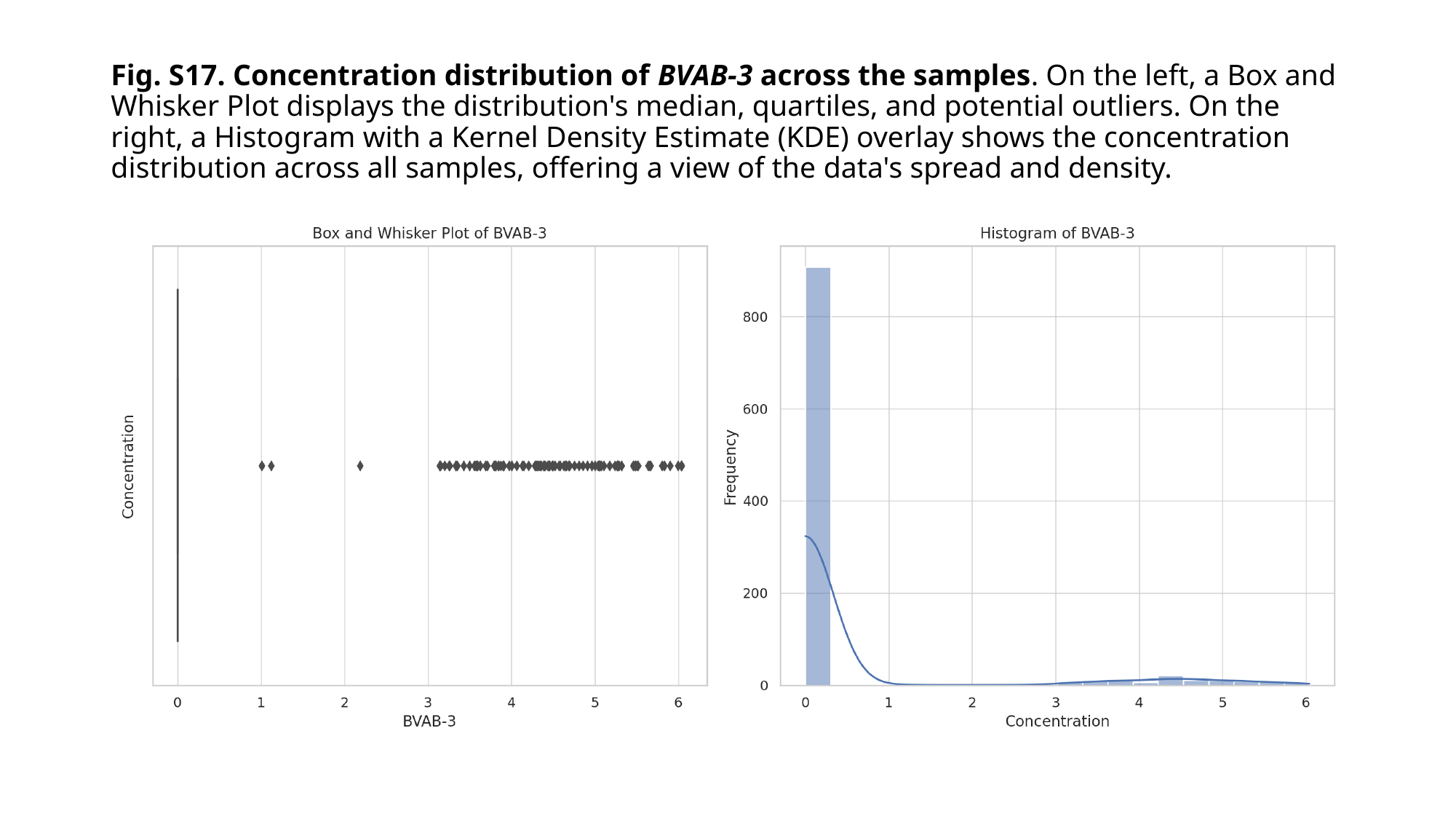

#### Fig. S17. Concentration distribution of BVAB-3 across the samples. On the left, a Box and Whisker Plot displays the distribution's median, quartiles, and potential outliers. On the right, a Histogram with a Kernel Density Estimate (KDE) overlay shows the concentration distribution across all samples, offering a view of the data's spread and density.

### Slide 19
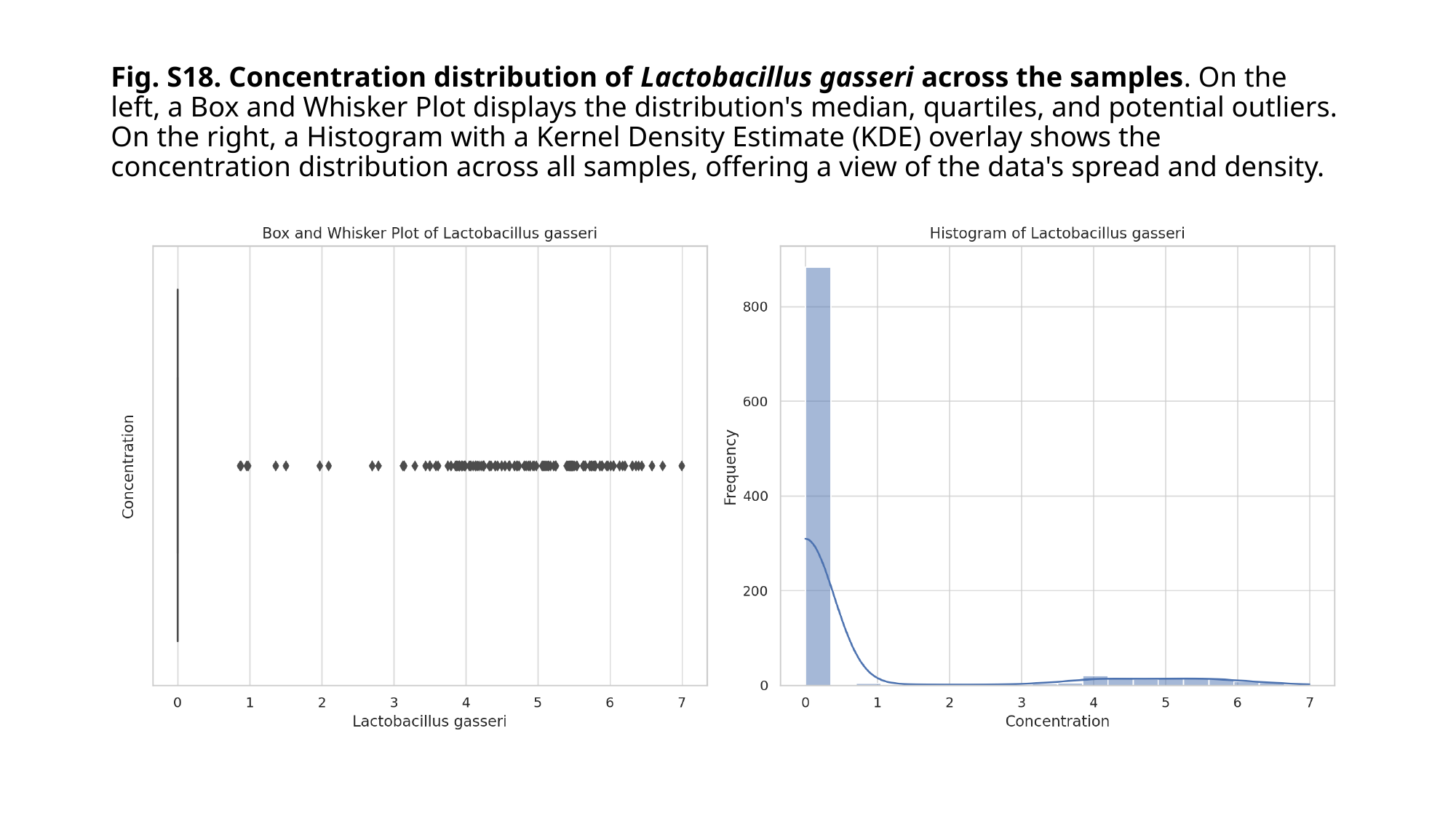

#### Fig. S18. Concentration distribution of Lactobacillus gasseri across the samples. On the left, a Box and Whisker Plot displays the distribution's median, quartiles, and potential outliers. On the right, a Histogram with a Kernel Density Estimate (KDE) overlay shows the concentration distribution across all samples, offering a view of the data's spread and density.

### Slide 20
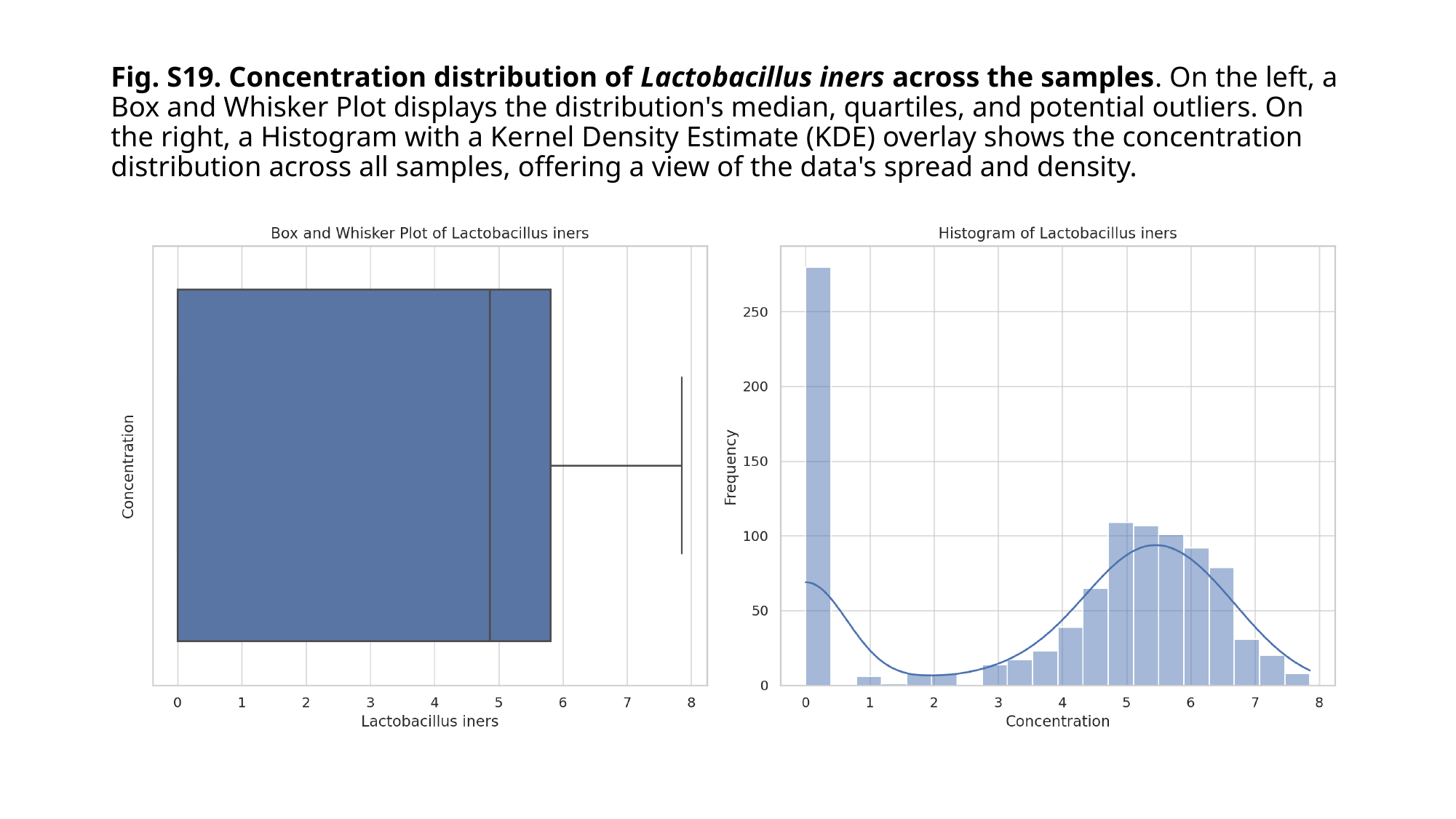

#### Fig. S19. Concentration distribution of Lactobacillus iners across the samples. On the left, a Box and Whisker Plot displays the distribution's median, quartiles, and potential outliers. On the right, a Histogram with a Kernel Density Estimate (KDE) overlay shows the concentration distribution across all samples, offering a view of the data's spread and density.

### Slide 21
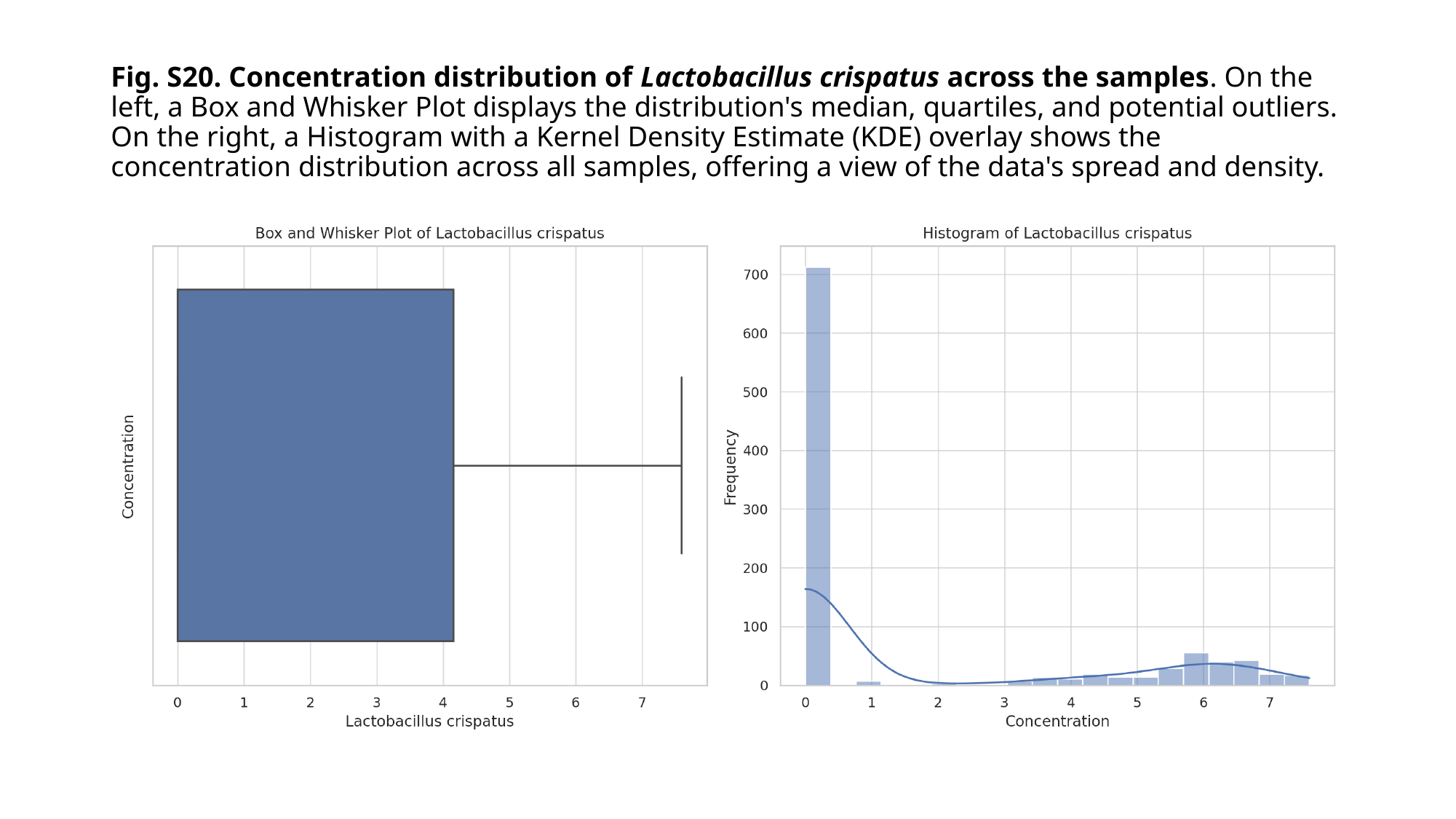

#### Fig. S20. Concentration distribution of Lactobacillus crispatus across the samples. On the left, a Box and Whisker Plot displays the distribution's median, quartiles, and potential outliers. On the right, a Histogram with a Kernel Density Estimate (KDE) overlay shows the concentration distribution across all samples, offering a view of the data's spread and density.

### Slide 22
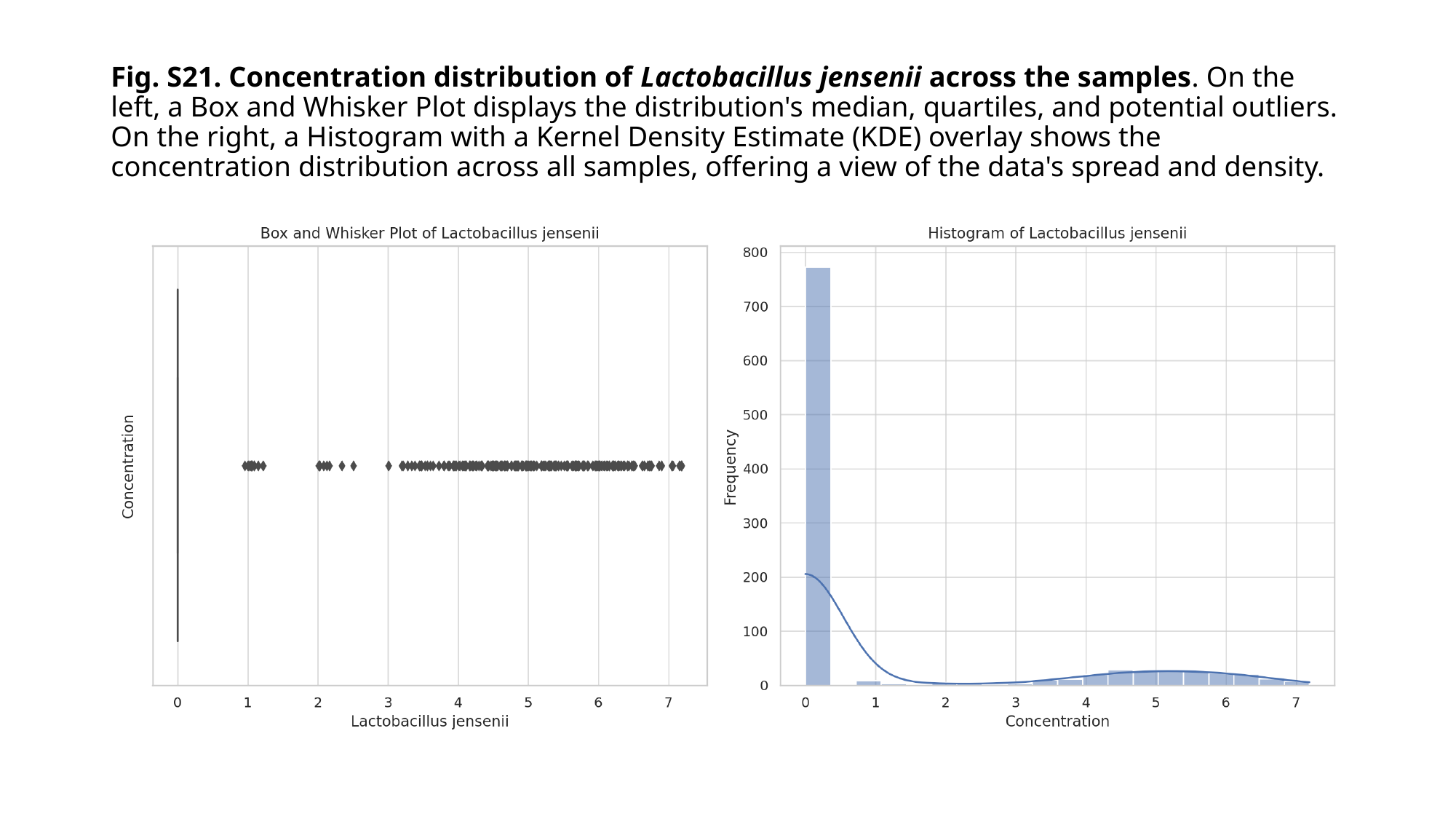

#### Fig. S21. Concentration distribution of Lactobacillus jensenii across the samples. On the left, a Box and Whisker Plot displays the distribution's median, quartiles, and potential outliers. On the right, a Histogram with a Kernel Density Estimate (KDE) overlay shows the concentration distribution across all samples, offering a view of the data's spread and density.

### Slide 23
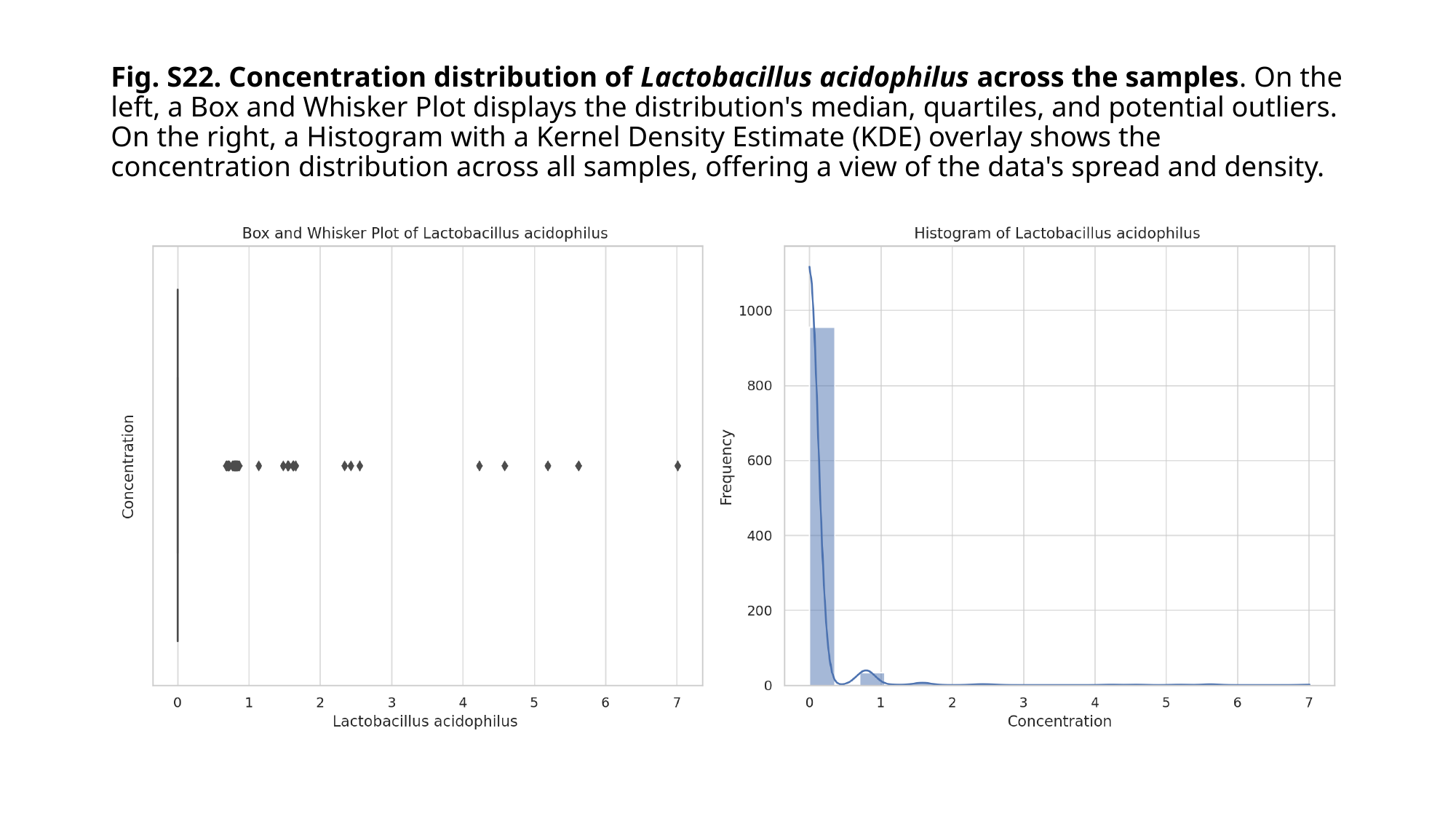

#### Fig. S22. Concentration distribution of Lactobacillus acidophilus across the samples. On the left, a Box and Whisker Plot displays the distribution's median, quartiles, and potential outliers. On the right, a Histogram with a Kernel Density Estimate (KDE) overlay shows the concentration distribution across all samples, offering a view of the data's spread and density.

### Slide 24
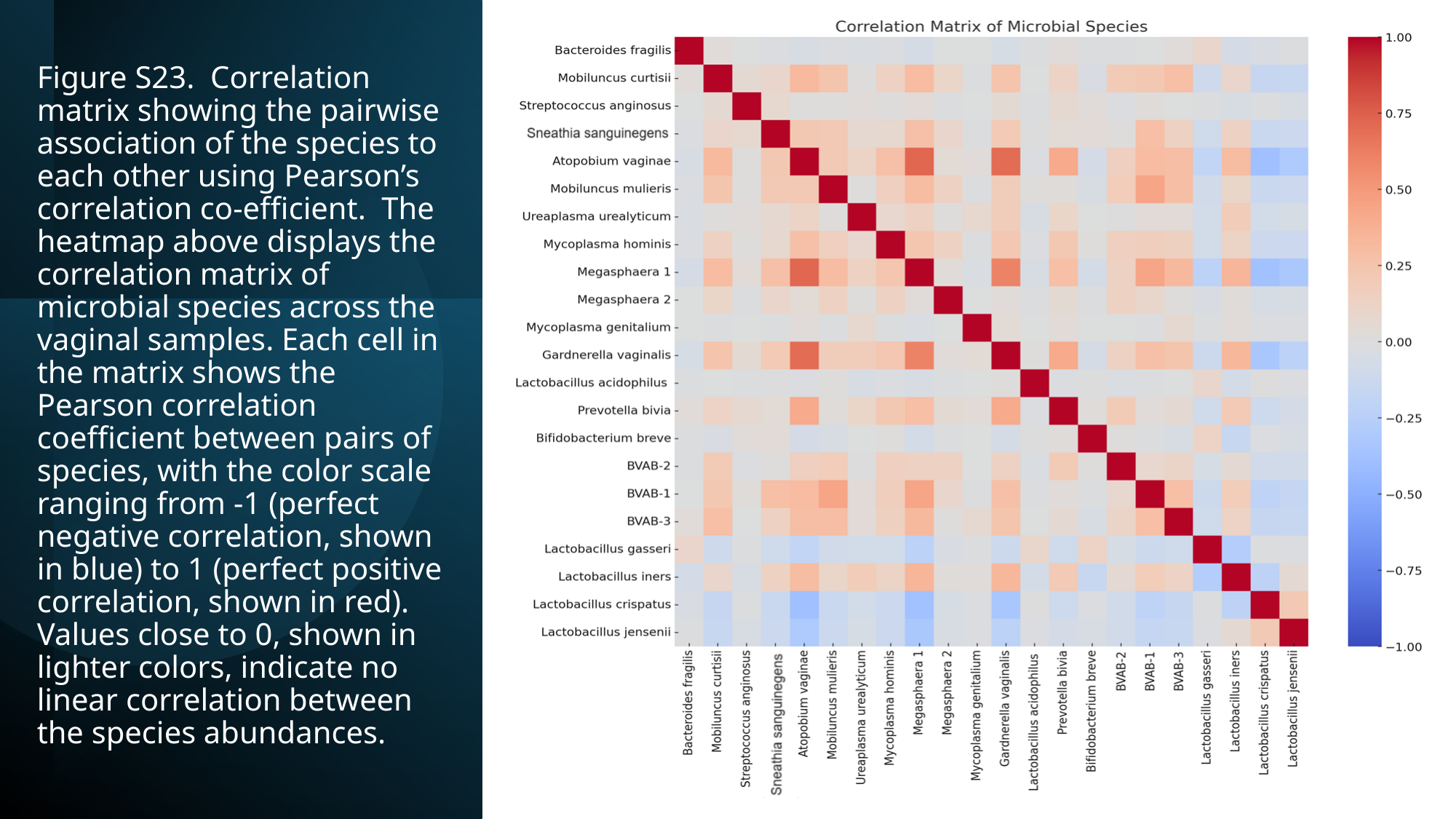

#### Figure S23. Correlation matrix showing the pairwise association of the species to each other using Pearson’s correlation co-efficient. The heatmap above displays the correlation matrix of microbial species across the vaginal samples. Each cell in the matrix shows the Pearson correlation coefficient between pairs of species, with the color scale ranging from -1 (perfect negative correlation, shown in blue) to 1 (perfect positive correlation, shown in red). Values close to 0, shown in lighter colors, indicate no linear correlation between the species abundances.

### Slide 25
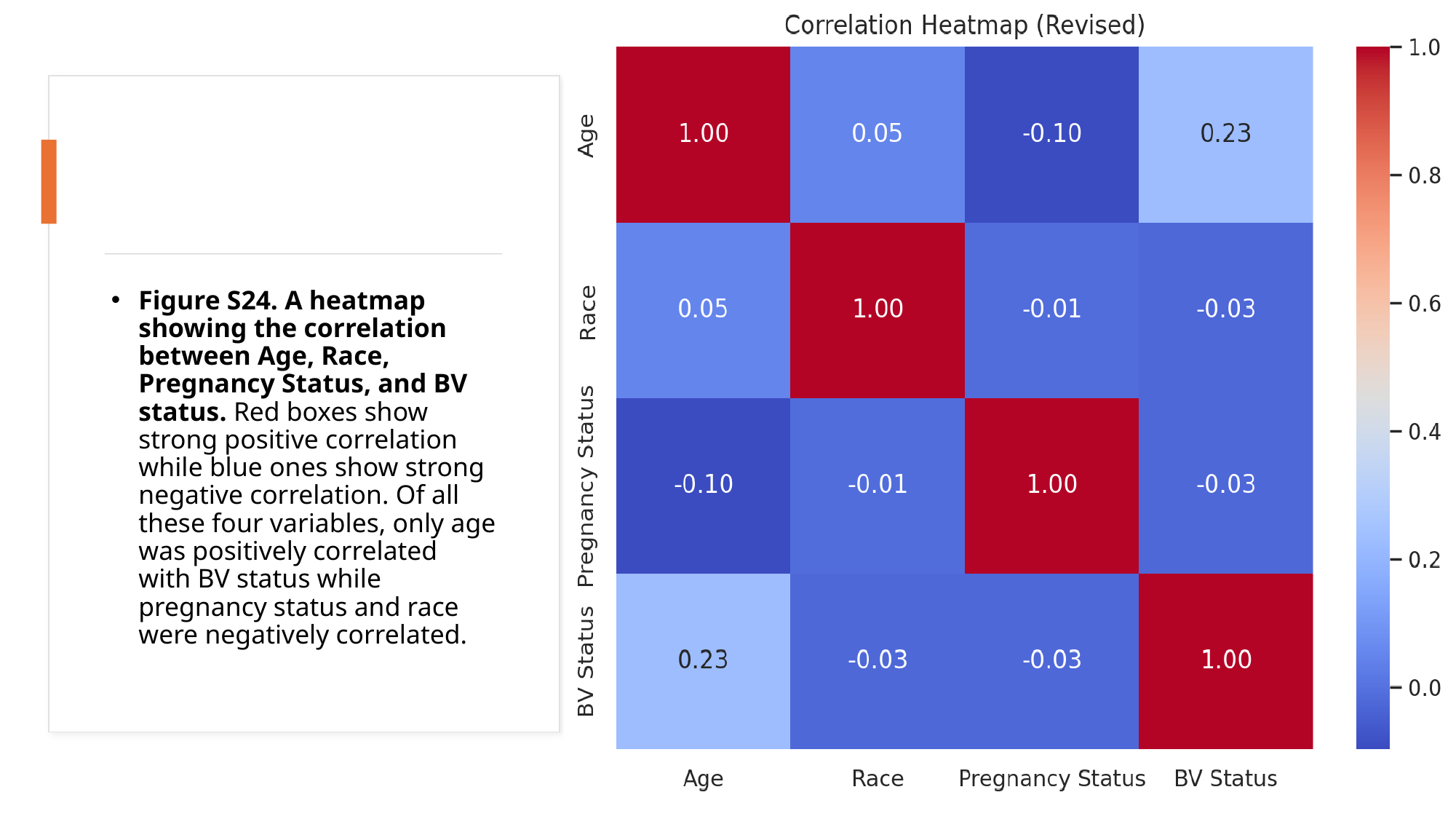

Figure S24. A heatmap showing the correlation between Age, Race, Pregnancy Status, and BV status. Red boxes show strong positive correlation while blue ones show strong negative correlation. Of all these four variables, only age was positively correlated with BV status while pregnancy status and race were negatively correlated.
